# AMP-activated protein kinase is necessary for Treg cell functional adaptation to microenvironmental stress

**DOI:** 10.1101/2023.11.29.568904

**Authors:** Manuel A. Torres Acosta, Nurbek Mambetsariev, Carla P. Reyes Flores, Kathryn A. Helmin, Qianli Liu, Anthony M. Joudi, Luisa Morales-Nebreda, Jonathan Gurkan, Kathleen Cheng, Hiam Abdala-Valencia, Samuel E. Weinberg, Benjamin D. Singer

## Abstract

CD4+FOXP3+ regulatory T (Treg) cells maintain self-tolerance, suppress the immune response to cancer, and protect against tissue injury in the lung and other organs. Treg cells require mitochondrial metabolism to exert their function, but how Treg cells adapt their metabolic programs to sustain and optimize their function during an immune response occurring in a metabolically stressed microenvironment remains unclear. Here, we tested whether Treg cells require the energy homeostasis-maintaining enzyme AMP-activated protein kinase (AMPK) to adapt to metabolically aberrant microenvironments caused by malignancy or lung injury, finding that AMPK is dispensable for Treg cell immune-homeostatic function but is necessary for full Treg cell function in B16 melanoma tumors and during acute lung injury caused by influenza virus pneumonia. AMPK-deficient Treg cells had lower mitochondrial mass and exhibited an impaired ability to maximize aerobic respiration. Mechanistically, we found that AMPK regulates DNA methyltransferase 1 to promote transcriptional programs associated with mitochondrial function in the tumor microenvironment. In the lung during viral pneumonia, we found that AMPK sustains metabolic homeostasis and mitochondrial activity. Induction of DNA hypomethylation was sufficient to rescue mitochondrial mass in AMPK-deficient Treg cells, linking DNA methylation with AMPK function and mitochondrial metabolism. These results define AMPK as a determinant of Treg cell adaptation to metabolic stress and offer potential therapeutic targets in cancer and tissue injury.

## Introduction

Regulatory T (Treg) cells are a subset of CD4+ T cells defined by the expression of the Forkhead box P3 (FOXP3) transcription factor that maintain self-tolerance via the suppression of self-reactive effector immune cells (*1, 2*). Treg cells also regulate immune responses to cancer and acute inflammatory processes such as infections and tissue injury (*3*). In the tumor microenvironment (TME), Treg cell-mediated immune suppression becomes maladaptive and dampens the anti-tumor immune response to promote tumor progression (*4, 5*). In contrast, during acute inflammation such as in viral pneumonia, Treg cells promote tissue protection and recovery by restraining inflammation and coordinating the repair of the injured lung parenchyma (*6, 7*). Treg cell suppressive function is regulated by cellular metabolism, and while Treg cells upregulate glucose consumption when proliferating, their suppressive function requires oxidative phosphorylation and is dependent on mitochondrial metabolism (*8-11*). The central role of cellular metabolism in determining Treg cell function has been well described in the TME, where Treg cells rewire their nutrient uptake to adapt to the metabolic aberrations of the microenvironment and thereby sustain their suppressive function (*12, 13*). Despite the known causal association between cellular metabolism and Treg cell function, how Treg cells sense microenvironmental changes and undergo metabolic adaptation during microenvironmental stress to optimize their suppressive function is unclear.

AMP-activated protein kinase (AMPK) is a heterotrimeric protein complex that serves as a master regulator of cellular metabolism (*14*). In settings of energetic stress, adenosine monophosphate (AMP) binds to AMPK and promotes its activation, priming the complex to phosphorylate downstream targets that mediate the restoration of energy homeostasis via one of two α catalytic subunits (AMPKα1, encoded by the *Prkaa1* gene or AMPKα2, encoded by the *Prkaa2* gene) (*15, 16*). Canonically, AMPK effects its energy-replenishing function through the phosphorylation of cytoplasmic factors; however, recent work *in vitro* supports an emerging role for AMPK as a regulator of epigenetic modifiers, including DNA methyltransferase 1 (DNMT1) (*17, 18*). Whether AMPK activates metabolic transcriptional programs via epigenetic mechanisms in immune cells *in vivo* remains unknown.

The two isoforms of the catalytic subunit of AMPK (AMPKα1/α2) are dispensable for *in vivo* Treg cell-mediated immune self-tolerance, but it is unclear whether Treg cells require AMPKα1/α2 to regulate acute immune responses in metabolically stressed microenvironments (*19-22*). Considering AMPK’s role in sustaining energy homeostasis via the potentiation of mitochondrial metabolism, and the necessity of oxidative phosphorylation (OXPHOS) for Treg cell suppressive function, we hypothesized that Treg cells require AMPK during states of metabolic stress to potentiate mitochondrial metabolism and thereby optimize Treg cell suppressive function. To test our hypothesis, we generated Treg cell-specific AMPKα1- and AMPKα2-deficient mice (*Prkaa1^fl/fl^Prkaa2^fl/fl^Foxp3^YFP-Cre^*, referred to here as *Prkaa1/2^fl/fl^Foxp3^YFP-Cre^* mice) and challenged them with either subcutaneous B16 mel anoma tumors or intra-tracheal inoculations of influenza A/WSN/33 H1N1 virus, both disease models whose outcomes are dependent on Treg cell function and whose microenvironments are burdened with metabolic derangements that challenge cellular metabolism (*4, 23*). We confirmed that AMPKα1/α2 are dispensable for Treg cell-mediated immune self-tolerance but found that *Prkaa1/2^fl/fl^Foxp3^YFP-Cre^*mice grew smaller tumors and experienced greater mortality and hypoxemia during influenza, with evidence of greater intra-tumoral and lung immune activation. Mechanistically, loss of AMPKα1/α2 in Treg cells resulted in accumulation of DNMT1 protein and promoter DNA hypermethylation at specific loci encoding metabolic genes, which were transcriptionally repressed. Consistent with this downregulation of metabolic gene expression, AMPKα1/α2-deficient Treg cells displayed impaired mitochondrial metabolism at homeostasis, in the TME, and in influenza virus-infected lungs. Pharmacologic-mediated induction of DNA hypomethylation rescued mitochondrial mass in AMPKα1/α2-deficient Treg cells, demonstrating that DNA methylation regulates Treg cell mitochondrial mass in an AMPK-dependent manner. In summary, our data indicates that AMPK is necessary to maintain epigenetic and metabolic programs that support optimal Treg cell suppressive function in metabolically stressed microenvironments such as the TME and the lung during viral pneumonia.

## Results

### AMPKα is dispensable for Treg cell suppressive function under homeostatic conditions

We confirmed loss of AMPKα1/α2 in CD4+FOXP3+ Treg cells (see **Supplemental Fig 1A** for gating strategy) isolated from *Prkaa1/2^fl/fl^Foxp3^YFP-Cre^* mice, which bred in approximately mendelian sex ratios (**Supplemental Fig 1B-E**). Consistent with previous reports (*19-22*), a tissue survey of spleen, thymus, and lungs did not reveal significant differences in CD8+ T cell infiltration between *Prkaa1/2^fl/fl^Foxp3^YFP-Cre^* and *Prkaa1/2^wt/wt^Foxp3^YFP-Cre^*(control) mice (**Fig 1A**). Supporting a lack of spontaneous inflammation resulting from Treg cell-specific loss of AMPKα1/α2, there were no significant differences between *Prkaa1/2^fl/fl^Foxp3^YFP-Cre^*and control mice in their spleen mass or the relative proportion of naïve (CD62L^Hi^CD44^Lo^) and effector (CD62L^Lo^CD44^Hi^) splenic conventional T (Tconv) cells (**Fig 1B-C**). Although the total number and proliferation rate of Treg cells was not significantly different between groups (**Fig 1D-E**), the splenic Treg cell compartment in *Prkaa1/2^fl/fl^Foxp3^YFP-Cre^*mice displayed a nominal yet statistically significant shift toward a central (CD62L^Hi^CD44^Lo^) Treg cell phenotype relative to control mice (**Fig 1F**). The *Foxp3^YFP-Cre^* allele used to drive *Foxp3*-dependent expression of Cre recombinase also drives expression of yellow fluorescent protein (YFP), which serves as a transcriptional reporter for the *Foxp3* locus. Treg cells from *Prkaa1/2^fl/fl^Foxp3^YFP-Cre^* mice showed similar expression of *Foxp3*-YFP to control mice, although we detected slightly lower FOXP3 protein in AMPKα1/α2-deficient splenic Treg cells measured by direct conjugated antibody staining (**Fig 1G-H**). AMPKα1/α2-deficient Treg cells displayed no significant differences in their ability to suppress responder CD4+ Tconv cell proliferation *in vitro* relative to controls (**Fig 1I**) and showed no significant differences in their surface membrane levels of markers traditionally correlated with Treg cell suppressive function (CD25, CTLA-4, PD-1, TIGIT, and ICOS, **Supplemental Fig 1F-J**) or their proliferation rate *in vitro* (**Supplemental Fig 1K**). Pharmacologic activation of AMPK promotes *Foxp3* expression *in vitro* (*24, 25*). To test whether AMPKα1/α2 are necessary for generation of induced (i)Treg cells, we subjected sorted CD4+Foxp3-Tconv cells from *Prkaa1/2^fl/fl^Foxp3^YFP-Cre^* and control mice to Treg cell-polarizing conditions for 5 days. We detected no significant difference in *Foxp3* induction efficiency between groups (**Supplemental Fig 1L**). We also performed unsupervised assessment of the metabolome of AMPKα1/α2-deficient and control splenic Treg cells using liquid chromatography tandem mass spectrometry (LC-MS) but found no significant differences between groups across the measured metabolites (258 annotated metabolites, **Supplemental Fig 1M** and **Supplemental File 1**). Finally, we assessed the transcriptional state of AMPKα1/α2-deficient and control splenic Treg cells at homeostasis via RNA-sequencing and identified 78 differentially expressed genes (DEGs) (**Fig 1J** and **Supplemental File 2**). Among the genes downregulated in AMPKα1/α2-deficient splenic Treg cells were components of the electron transport chain (*mt-Nd2* and *mt-Co1*) and heat shock proteins (*Hspa1a*, *Hspa1b*, and *Hspa8*), consistent with AMPK’s positive regulation of mitochondrial metabolism and the cellular stress response. Genes upregulated in AMPKα1/α2-deficient splenic Treg cells included cytokines and transcription factors associated with effector T cell function (*Tnf*, *Nfkbid*, and *Rora*) and regulators of one-carbon metabolism (*Mthfr*). Gene set enrichment analysis (GSEA) demonstrated downregulation of genes associated with Treg cell identity and function (*26*) (**Fig 1K**), suggesting that although AMPKα1/α2 are dispensable for Treg cell-mediated immune self-tolerance during development and homeostasis, AMPKα1/α2-deficient Treg cells may suffer functional impairment in settings requiring enhanced suppressive function, such as the TME.

**Figure 1.**
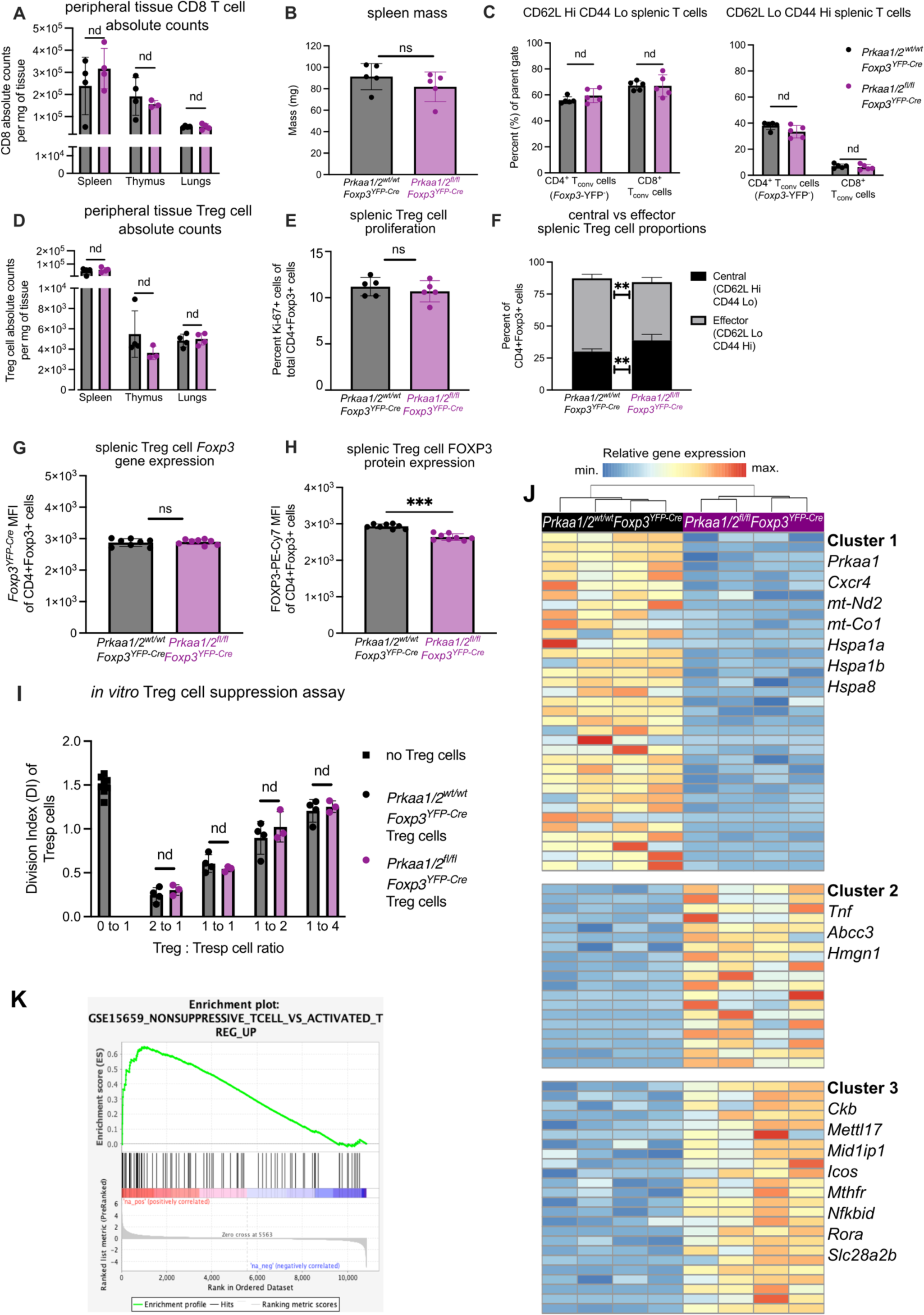
AMPKα1 and AMPKα2 are dispensable for Treg cell-mediated immune self-tolerance and Treg cell suppressive function at homeostasis. (**A**) CD8+ conventional T (Tconv) cell absolute counts per milligram (mg) of *Prkaa1/2^wt/wt^Foxp3^YFP-Cre^*(control) and *Prkaa1/2^fl/fl^Foxp3^YFP-Cre^* mouse spleen (*n*=4 control, *n*=4 *Prkaa1/2^fl/fl^Foxp3^YFP-Cre^*), thymus (*n*=4 control, *n=*3 *Prkaa1/2^fl/fl^Foxp3^YFP-Cre^*), and lung (*n*=4 control, *n*=4 *Prkaa1/2^fl/fl^Foxp3^YFP-Cre^*). (**B**) Spleen mass of 8–12-week-old control (*n*=5) and *Prkaa1/2^fl/fl^Foxp3^YFP-Cre^*(*n*=5) mice. (**C**) Frequency of naive (CD62L^Hi^CD44^Lo^) and effector (CD62^Lo^CD44^Hi^) splenic CD8+ and CD4+ Tconv cells out of total CD8+ and CD4+ cells, respectively (*n*=5 control, *n*=5 *Prkaa1/2^fl/fl^Foxp3^YFP-Cre^*). (**D**) CD4+Foxp3+ cell absolute counts per mg of control and *Prkaa1/2^fl/fl^Foxp3^YFP-Cre^* mouse spleen (*n*=4 control, *n*=4 *Prkaa1/2^fl/fl^Foxp3^YFP-Cre^*), thymus (*n*=4 control, *n*=3 *Prkaa1/2^fl/fl^Foxp3^YFP-Cre^*), and lung (*n*=4 control, *n*=4 *Prkaa1/2^fl/fl^Foxp3^YFP-Cre^*). (**E**) Frequency of Ki- 67+CD4+Foxp3+ cells out of total CD4+Foxp3+ splenocytes (*n*=5 control, *n*=5 *Prkaa1/2^fl/fl^Foxp3^YFP-Cre^*). (**F**) Frequency of central (CD62L^Hi^CD44^Lo^) and effector (CD62^Lo^CD44^Hi^) CD4+Foxp3+cells out of total CD4+Foxp3+ splenocytes (*n*=5 control, *n*=5 *Prkaa1/2^fl/fl^Foxp3^YFP-Cre^*). (**G-H**) *Foxp3*-YFP (**G**) and FOXP3-PE-Cy7 (**H**) mean fluorescence intensity (MFI) of CD4+Foxp3+ splenocytes (*n*=8 control, *n*=8 *Prkaa1/2^fl/fl^Foxp3^YFP-Cre^*). (**I**) Division index of CD4+Foxp3-splenic responder T (Tresp) cells co-cultured for 72 hours with varying ratios of CD4+Foxp3+ splenocytes (*n*=4 control, *n*=3 *Prkaa1/2^fl/fl^Foxp3^YFP-Cre^*). (**J**) *K*-means clustering of 87 significant differentially expressed genes (FDR *q*-value < 0.05) identified between CD4+Foxp3+ cells sorted from spleens of control (*n*=4) and *Prkaa1/2^fl/fl^Foxp3^YFP-Cre^* (*n*=4) mice with *k*=3 and scaled as *Z*-scores across rows. (**K**) Enrichment plot of the GSE15659_NONSUPPRESSIVE_TCELL_VS_ ACTIVATED_TREG_UP geneset (consists of genes down-regulated in comparison of resting Treg versus non-suppressive T cells) generated through gene set enrichment analysis (GSEA) preranked testing of the expressed genes of *Prkaa1/2^fl/fl^Foxp3^YFP-Cre^*and control splenic Treg cells identified by RNA-sequencing. ** *p* or *q* < 0.01; *** *p* or *q* < 0.001; nd, no discovery, ns, not significant according to Mann-Whitney *U* test (**B, E, G, H**) with two-stage linear step-up procedure of Benjamini, Krieger, and Yekutieli with Q = 5% (**A, C, D, F, I**). Summary plots show all data points with mean and SD.

### AMPKα promotes Treg cell suppressive function in the tumor microenvironment

While we did not detect *Prkaa2* expression in splenic and lymph node Treg cells of mice bearing subcutaneous B16 melanoma tumor grafts, we found that *Prkaa1/2^wt/wt^Foxp3^YFP-Cre^*(control) Treg cells upregulated the expression of both *Prkaa1* and *Prkaa2* in the TME (**Supplemental Fig 2A-B**). Hence, to determine whether AMPKα1/α2-deficient Treg cells are functionally impaired in the TME, we challenged *Prkaa1/2^fl/fl^Foxp3^YFP-Cre^* and control mice with B16 melanoma tumors, finding that *Prkaa1/2^fl/fl^Foxp3^YFP-Cre^* mice experienced lower tumor burden over time (**Fig 2A**). Tumors of *Prkaa1/2^fl/fl^Foxp3^YFP-Cre^* mice had significantly higher CD8-to-Treg cell ratios relative to controls at day 15 post-engraftment (**Fig 2B**), consistent with a loss of Treg cell suppressive function in the TME. We did not find significant differences in the intra-tumor proportion of naïve, central memory, or effector Tconv cell subsets between groups at day 15 post-engraftment (**Supplemental Fig 2C-G**). There were also no significant differences between the Treg cells of *Prkaa1/2^fl/fl^Foxp3^YFP-Cre^* and control mice in their abundance out of all CD4+ cells, their proportion of central versus effector subsets, their proliferation, or their *Foxp3* gene and FOXP3 protein expression (**Supplemental Fig 2H-M**). Similar to the spleen at homeostasis, most traditional surface markers of Treg cell suppressive function were not significantly different between tumor-infiltrating AMPKα1/α2-deficient and control Treg cells (**Supplemental Fig 2N-R**). We leveraged RNA-sequencing to profile the transcriptional state of Treg cells sorted from the tumors of *Prkaa1/2^fl/fl^Foxp3^YFP-Cre^* and control mice at day 15 post-engraftment and identified 752 DEGs (**Fig 2C**). Unsupervised clustering revealed that the two largest groups of DEGs (Clusters 1 and 2) were downregulated in AMPKα1/α2-deficient cells (**Fig 2D** and **Supplemental File 3**). The *Ppargc1a* gene, encoding a master regulator of mitochondrial biogenesis, PGC-1α, was present in Cluster 1 (**Fig 2E**). Accordingly, functional enrichment analysis demonstrated that Cluster 1 genes are involved in cellular metabolism and include Gene Ontology (GO) terms relating to cellular response to stress and mitochondrial metabolism; cluster 2 genes are involved in immune effector cell programs and in epigenetic regulation of transcription (**Fig 2F**). Analysis of cluster 3 genes, which were upregulated in AMPKα1/α2-deficient Treg cells, linked this cluster to a broad set of cellular functions including negative regulation of transcription. Gene Set Enrichment Analysis revealed a positive enrichment of genes associated with allograft rejection and interferon gamma signaling as well as a negative enrichment of genes associated with angiogenesis in tumor-infiltrating AMPKα1/α2-deficient Treg cells (**Fig 2G-I**), consistent with loss of Treg cell function in the TME. In addition, tumor-infiltrating AMPKα1/α2-deficient Treg cells also showed transcriptional signatures associated with downregulated response to hypoxia, glycolysis, and cholesterol homeostasis (**Fig 2J-L**), suggestive of failed metabolic adaptation in the TME. Collectively, these data suggest that tumor-infiltrating AMPKα1/α2-deficient Treg cells fail to upregulate metabolic and effector immune cell transcriptional programs, thereby compromising suppressive function in the TME.

**Figure 2.**
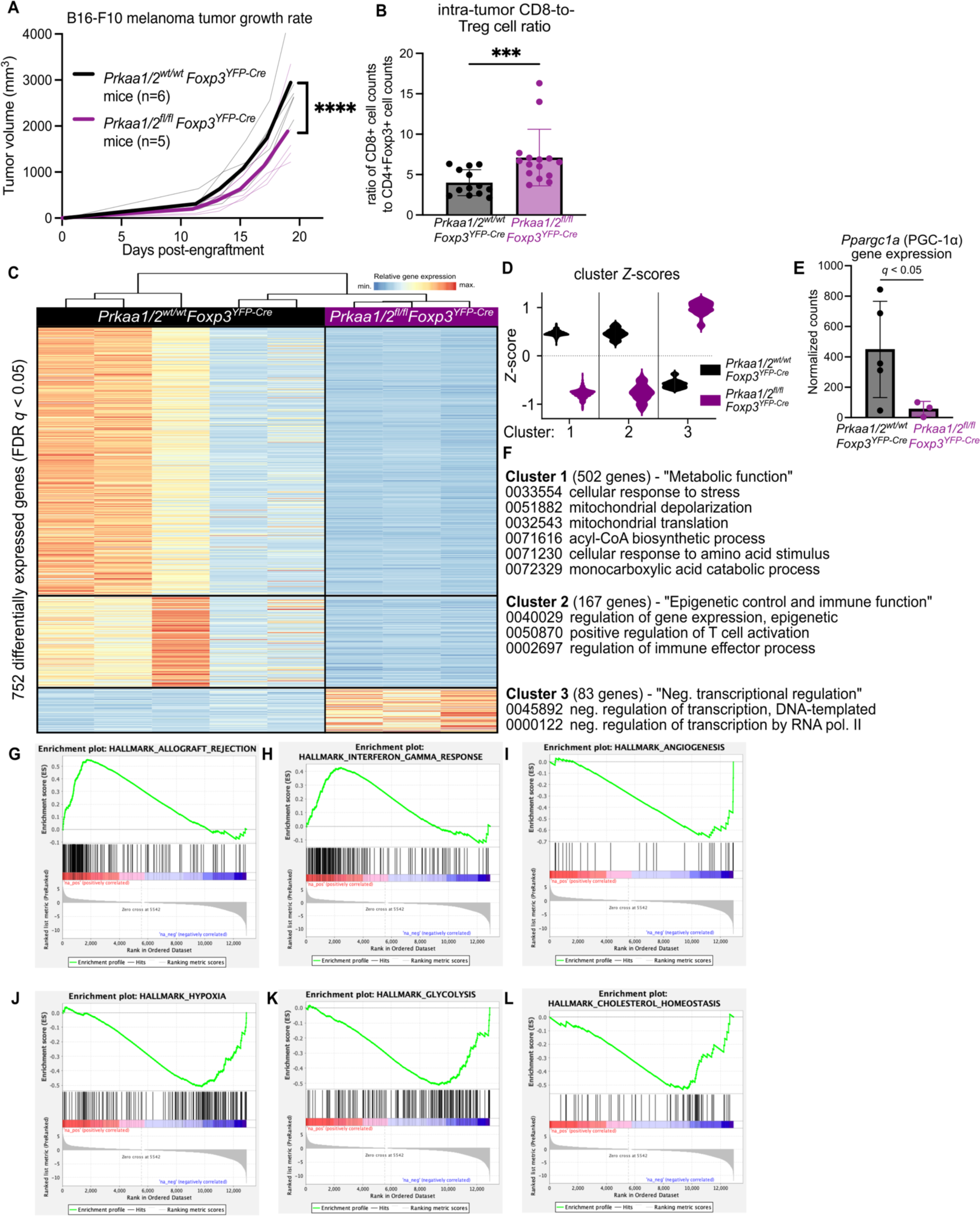
AMPKα1/α2 loss is sufficient to impair Treg cell suppressive function in the tumor microenvironment. (**A**) Growth of B16 melanoma tumors in *Prkaa1/2^wt/wt^Foxp3^YFP-Cre^*(control, *n*=6) and *Prkaa1/2^fl/fl^Foxp3^YFP-Cre^*(*n*=5) mice. (**B**) Ratio of live CD8+ cell counts to live CD4+Foxp3+ cell counts in single cell suspensions of B16 melanoma tumors harvested from the flanks of control (*n*=13) and *Prkaa1/2^fl/fl^Foxp3^YFP-Cre^* (*n*=15) mice at day 15 post-engraftment. (**C**) *K*-means clustering of 752 significant differentially expressed genes (FDR *q*-value < 0.05) identified between CD4+Foxp3+ cells sorted from B16 melanoma tumors of control (*n*=5) and *Prkaa1/2^fl/fl^Foxp3^YFP-Cre^* (*n*=3) mice with *k*=3 and scaled as *Z*-scores across rows. (**D**) Average z-scores for the three clusters shown in (**C**). (**E**) Normalized counts of *Ppargc1a* reads measured by RNA-sequencing (*n*=5 control, *n*=3 *Prkaa1/2^fl/fl^Foxp3^YFP-Cre^*). (**F**) Selection of top gene ontology (GO) processes derived from Clusters 1, 2, and 3 (all with FDR *q* < 0.05). (**G**-**L**) Enrichment plots (*p* < 0.05, FDR < 0.25) of the HALLMARK_ALLOGRAFT_REJECTION gene set (G), HALLMARK_INTERFERON_GAMMA_ RESPONSE gene set (H), HALLMARK_ANGIOGENESIS gene set (I), HALLMARK_HYPOXIA gene set (J), HALLMARK_GLYCOLYSIS gene set (K), and HALLMARK_CHOLESTEROL HOMEOSTASIS gene set (L). Enrichment plots were generated through GSEA preranked testing of the expressed genes of tumor-infiltrating *Prkaa1/2^fl/fl^Foxp3^YFP-Cre^* and control Treg cells identified by RNA-sequencing. **** *p* < 0.0001 according to 2-way ANOVA with two-stage linear step-up procedure of Benjamini, Krieger, and Yekutieli with Q = 5% (**A**). *** *p* <0.001 according to Mann Whitney U test (**B**). Summary plots show all data points with mean and SD.

### AMPKα2 contributes to the regulation of Treg cell suppressive function in the tumor microenvironment

Previous studies that evaluated the requirement of AMPK for Treg cell-mediated suppression of the anti-tumor immune response leveraged mouse models of Treg cell-specific AMPKα1-conditional knockout mice (*Prkaa1^fl/fl^Foxp3^YFP-Cre^*), and the results have been conflicting on whether loss of AMPKα1 potentiates or compromises Treg cell function in the TME (*27, 28*). To test the significance of our finding that Treg cells upregulate *Prkaa2* in the TME (see **Supplemental Fig 2B**), we generated Treg cell-specific AMPKα1-deficient (*Prkaa1^fl/fl^Foxp3^YFP-Cre^*) and AMPKα2-deficient (*Prkaa2^fl/fl^Foxp3^YFP-Cre^*) mice and evaluated their response to B16 melanoma tumors. We observed significantly smaller tumors in Treg cell-specific AMPKα1-deficient mice relative to controls, while those with AMPKα2-deficient Treg cells exhibited significantly greater tumor volume over time through day 15 post-tumor engraftment relative to mice bearing control, AMPKα1-, and AMPKα1/α2-deficient Treg cells (**Supplemental Fig 3A-B**). Consistent with this finding, *Prkaa1^fl/fl^Foxp3^YFP-Cre^* mice had higher CD8-to-Treg cell ratios compared with *Prkaa2^fl/fl^Foxp3^YFP-Cre^* and control mice (**Supplemental Fig 3C**). Tumors of *Prkaa2^fl/fl^Foxp3^YFP-Cre^*mice also exhibited a shift in their CD8+ Tconv cell compartment toward a central memory (CD62L^Hi^CD44^Hi^) phenotype (**Supplemental Fig 3D-F**) and a decreased proportion of effector CD4+ Tconv cells relative to *Prkaa1^fl/fl^Foxp3^YFP-Cre^*and control mice (**Supplemental Fig 3G-H**). *Prkaa1^fl/fl^Foxp3^YFP-Cre^* mice had a significantly higher proportion of effector CD8+ T cells (CD62L^Lo^CD44^Hi^) relative to *Prkaa2^fl/fl^Foxp3^YFP-Cre^* mice and had the lowest proportion of Treg cells out of the CD4+ T cell pool in their tumors (**Supplemental Fig 3I**). Nevertheless, we found no significant differences between groups in the proliferation rate of tumor-infiltrating Treg cells (**Supplemental Fig 3J**), their FOXP3 and CD25 protein levels measured by flow cytometry (**Supplemental Fig 3K-L**), and the relative proportion of central versus effector Treg cell subsets (**Supplemental Fig 3M**). When assessing the status of markers of Treg cell suppressive function, we found that loss of AMPKα1 or AMPKα2 had opposing effects on tumor-infiltrating Treg cell PD-1 expression, with loss of AMPKα1 leading to lower levels and loss of AMPKα2 leading to higher levels (**Supplemental Fig 3N**). Collectively, these data indicate that loss of AMPKα1-occupied AMPK complexes is sufficient to impair Treg cell suppressive function, while loss of AMPKα2-occupied complexes is sufficient to potentiate Treg cell suppressive function.

### The metabolic landscape of the virus-injured lung largely resembles the TME in its metabolite abundance; however, they differ in the abundance of key carbon sources

Treg cells must adapt their metabolism to function in the metabolically deranged microenvironment of the TME (*29*), but it remains undetermined whether other inflammatory microenvironments where Treg cells have critical functions, such as a virally infected lung (*30*), exhibit similar metabolic aberrations, and thereby present similar environmental stress to Treg cells. To compare the metabolic changes that occur in the TME and the virus-injured lung, we collected interstitial fluid (IF) extracted from mouse lungs 10 days after infection with influenza virus (peak injury), B16 melanoma tumors 15 days after engraftment, and paired plasma samples. We then measured hydrophilic metabolite abundance via LC-MS. Principal component analysis of the LC-MS data (303 annotated metabolites) revealed that the first principal component (PC1), which represents 57.9% of the variance in the dataset, captured the variance due to differences in metabolite abundance between lung and tumor IF relative to plasma (**Fig 3A-B** and **Supplemental File 4**). Key metabolites that contributed to PC1 include 2-hydroxyglutarate and lactate, which are overrepresented in tumor and flu IF relative to plasma (**Fig 3C-D**) and suggest a state of reduced mitochondrial electron transport chain activity in these disease microenvironments (*29*). The second principal component (PC2; 14.5% of the variance) was due to differences in metabolite abundance between influenza virus-infected lung and tumor IF. Interestingly, key carbon sources such as glucose and glutamine were more abundant in the infected flu IF and less abundant in tumor IF relative to plasma (**Fig 3E-F**). Overrepresentation analysis of the significant differentially represented metabolites in tumor IF relative to plasma (**Fig 3G** and **Supplemental Fig 4A-B**) and flu IF during viral pneumonia relative to plasma (**Fig 3H** and **Supplemental Fig 4C-D**) revealed that the TME and the lung during viral pneumonia undergo significant changes in similar metabolic pathways related to amino acid metabolism (**Fig 3I**). Nevertheless, direct comparison of tumor IF and flu IF metabolites revealed disease state-specific metabolite signatures, including an enrichment of metabolites related to tryptophan, cysteine, and methionine metabolism in flu IF (**Supplemental Fig 5A-D**). These data suggest that Treg cells and other immune cells may experience shared metabolic challenges in the TME and the injured lung during viral pneumonia but may use different carbon sources in these microenvironments.

**Figure 3.**
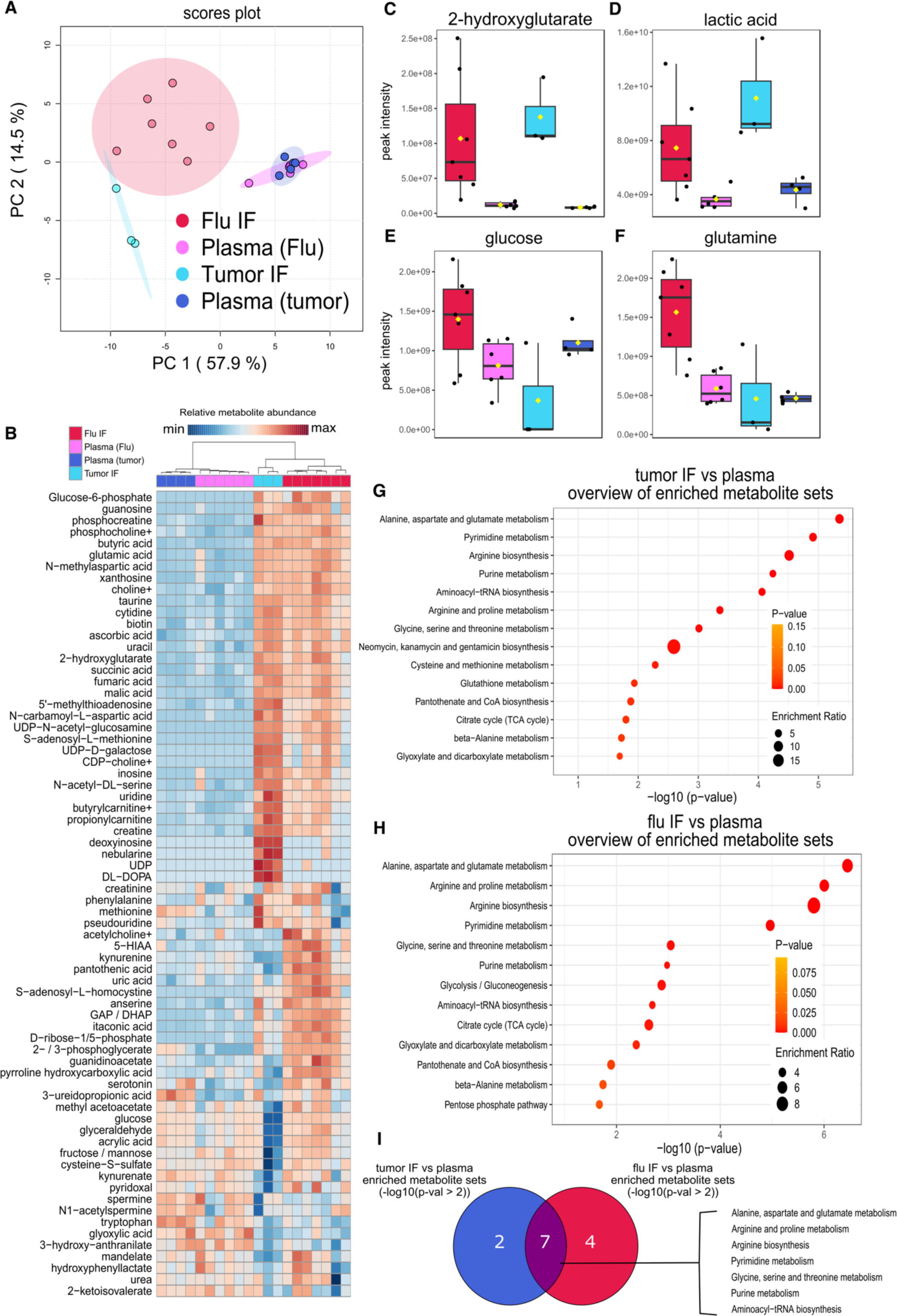
The metabolic landscape of the influenza virus-injured lung largely resembles the tumor microenvironment in its metabolite abundance; however, they differ in the abundance of key carbon sources. (**A**) Principal component (PC) analysis of the peak intensities of metabolites identified via liquid chromatography tandem mass spectrometry (LC-MS) from B16 melanoma tumor (*n*=3) and influenza virus-infected lung (flu, *n*=7) interstitial fluid (IF) and paired plasma (*n*=4 tumor, *n*=6 flu) from the same animals. (**B**) Heatmap of the 70 most differentially represented metabolites in plasma, tumor IF, and flu IF according to one-way ANOVA (*p*-val < 0.1). (**C-F**) Abundance of key significant differentially represented metabolites: 2-hydroxyglutarate (C), lactic acid (D), glucose (E), and glutamine (F). (**G**) Results from overrepresentation analysis of the significant (*p* < 0.1) differentially represented metabolites between tumor IF and plasma. (**H**) Results from overrepresentation analysis of the significant (*p* < 0.1) differentially represented metabolites between flu IF and plasma. (**I**) Overlap in significantly (*p* < 0.01) enriched metabolite sets between tumor IF vs plasma comparison and flu IF vs plasma comparison according to overrepresentation analysis of flu IF versus plasma and tumor IF versus plasma.

### AMPKα promotes Treg cell tissue-protective function during lung injury from viral pneumonia

Treg cells provide tissue protection following acute lung injury due to influenza virus infection and other causes of lung pathology and are necessary for resolution of inflammation and repair of lung injury during recovery (*30-33*). AMPKα1 is required for bulk CD4+ T cell expansion in the lung during viral pneumonia (*34*), but whether AMPKα1/α2 are necessary for Treg cell function in this context is unknown. We found that AMPKα1/α2-sufficient Treg cells express both *Prkaa1* and *Prkaa2* in lymph nodes and the lung during viral pneumonia (**Supplemental Fig 6A-B**) and that steady-state splenic AMPKα1/α2-deficient Treg cells have downregulated expression of genes activated during influenza A virus infection (*35*) (**Fig 4A**). Considering this transcriptional signature, our findings in the B16 melanoma model, and the similarity in interstitial fluid metabolite abundance between flu IF during viral pneumonia and tumor IF relative to plasma, we hypothesized that Treg cell-specific loss of AMPKα1/α2 would compromise protection from severe viral pneumonia. To test our hypothesis, we challenged *Prkaa1/2^fl/fl^Foxp3^YFP-Cre^*and control mice with intra-tracheal inoculations of influenza virus. *Prkaa1/2^fl/fl^Foxp3^YFP-Cre^*mice experienced higher mortality from a sublethal influenza A virus dose, greater weight loss throughout the disease course, and worsened hypoxemia (**Fig 4B-D**), consistent with a loss of Treg cell function. Accordingly, we detected a significant increase in the absolute number of lung CD45+ and CD8+ Tconv cells in *Prkaa1/2^fl/fl^Foxp3^YFP-Cre^* mice relative to controls at day 10 post-influenza virus inoculation (**Fig 4E-F**); lung Treg and CD4+ Tconv cell absolute counts were not significantly different across groups (**Fig 4G-H**). The CD8+ Tconv cell compartment displayed a nominal shift away from an effector phenotype in the lungs of *Prkaa1/2^fl/fl^Foxp3^YFP-Cre^*mice, but no significant differences were detected in the proportion of other lung CD8+ and CD4+ T cell subsets, including Treg cells (**Supplemental Fig 6C-J**). Interestingly, AMPKα1/α2-deficient Treg cells were less proliferative in the lung at day 10 post-influenza virus inoculation according to Ki-67 expression relative to control Treg cells (**Supplemental Fig K**). Nevertheless, like in our malignancy model, AMPKα1/α2 deficiency did not alter Treg cell *Foxp3*/FOXP3 gene/protein expression or the levels of markers associated with Treg cell suppressive function (**Supplemental Fig 6L-R**). As AMPK is a known regulator of cellular metabolism, we assessed the metabolic state of AMPKα1/α2-deficient lung Treg cells during viral pneumonia by performing LC-MS on AMPKα1/α2-sufficient and -deficient Treg cells sorted from lungs at day 10 post-influenza virus inoculation. 159 metabolites were annotated (**Fig 4I-J** and **Supplemental File 5**), revealing an enrichment of pyruvate and lactate in AMPKα1/α2-deficient lung Treg cells (**Fig 4K-L**), metabolites that are upstream of the tricarboxylic acid (TCA) cycle, suggestive of altered mitochondrial metabolism. We also detected depletion of glutathione (GSH), a key antioxidant, in AMPKα1/α2-deficient lung Treg cells (**Fig 4M**). Overrepresentation analysis of the significant differentially represented features revealed an overrepresentation of metabolites relating to glycine, serine, and threonine metabolism, glutathione metabolism, and pyruvate metabolism in AMPKα1/α2- deficient Treg cells (**Fig 4N**). Considering the absence of significant differences in metabolite abundance at homeostasis (see **Supplemental Fig 1M**), these data suggest AMPK is necessary for Treg cell metabolic adaptation and function during influenza virus-induced lung injury.

**Figure 4.**
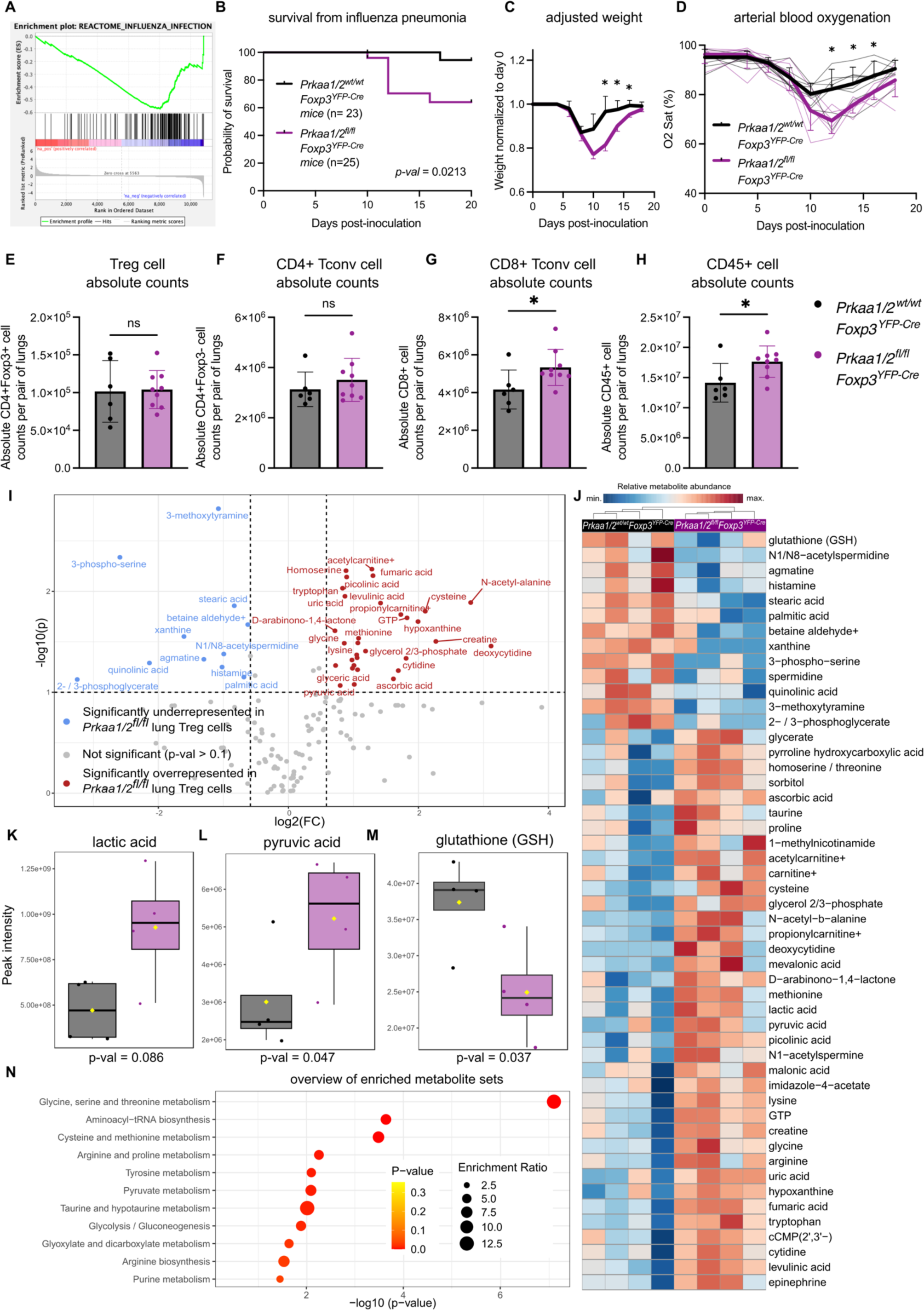
AMPKα1/α2 are necessary for optimal Treg cell function in the lung during influenza pneumonia. (**A**) Enrichment plot of the REACTOME_INFLUENZA_INFECTION geneset (*p* < 0.05, FDR < 0.25) generated through GSEA preranked testing of the expressed genes of *Prkaa1/2^wt/wt^Foxp3^YFP-Cre^*(control) and *Prkaa1/2^fl/fl^Foxp3^YFP-Cre^* CD4+Foxp3+ splenocytes identified by RNA-sequencing shown in Fig 1J. (**B**) Survival of control (*n*=23) and *Prkaa1/2^fl/fl^Foxp3^YFP-Cre^* (*n*=25) mice following intra-tracheal inoculation of 12.5 plaque forming units (PFUs) of influenza A/WSN/33 H1N1 (influenza) virus. (**C**-**D**) Weight (C), and arterial oxyhemoglobin saturation (D) over time of control (*n*=6) and *Prkaa1/2^fl/fl^Foxp3^YFP-Cre^* (*n*=8) mice following intra-tracheal inoculation of 12.5 PFUs of influenza virus. (**E-H**) Absolute counts of CD4+Foxp3+ cells (E), CD4+Foxp3-cells (F), CD8+ cells (G), and CD45+ cells (H) per pair of lungs in control (*n*=6) and *Prkaa1/2^fl/fl^Foxp3^YFP-Cre^* (*n*=9) mice at day 10 post-influenza virus inoculation. (**I**) Volcano plot of abundance of metabolites detected in control (*n*=4) and *Prkaa1/2^fl/fl^Foxp3^YFP-Cre^*(*n*=4) Treg cells sorted from lungs at day 10 post-influenza virus-inoculation. (**J**) Heatmap of top 50 differentially represented metabolites between control (*n*=4) and *Prkaa1/2^fl/fl^Foxp3^YFP-Cre^*(*n*=4) Treg cells sorted from lungs at day 10 post-influenza virus inoculation. (**K**-**M**) Peak intensities measured for lactic acid (K), pyruvic acid (L), and glutathione GSH (M) in Treg cells from the lungs of control (*n*=4) and *Prkaa1/2^fl/fl^Foxp3^YFP-Cre^* (*n*=4) mice at day 10 post-influenza virus-inoculation. (**N**) Results of overrepresentation analysis from the significant (*p* < 0.1, log2FC ≥ 1.5 or ≤ −1.5) differentially represented metabolites identified in (I). Survival curve (**B**) *p* was determined using log-rank (Mantel-Cox) test. * *q* < 0.05 according to two-way ANOVA with two-stage linear step-up procedure of Benjamini, Krieger, and Yekutieli with Q = 5% (**C**-**D**). * *p* < 0.05, ns not significant according to Mann-Whitney *U* test (**E**-**H**). Summary plots show all data points with mean and SD.

### AMPKα is necessary for maximal mitochondrial function in Treg cells

We demonstrated that mitochondrial metabolism, specifically activity of the electron transport chain, is a key determinant of Treg cell suppressive function (*10*). To test whether loss of AMPK compromises Treg cell mitochondrial function, we assessed the metabolic status of AMPKα1/α2-deficient and control Treg cells with a metabolic flux assay, finding that AMPKα1/α2-deficient Treg cells have comparable basal oxygen consumption rates (OCR) but significantly lower maximum OCR relative to control Treg cells (**Fig 5A-C**). In fact, AMPKα1/α2-deficient Treg cells were unable to augment their OCR above baseline when challenged with the mitochondrial uncoupling agent carbonyl cyanide m-chlorophenylhydrazone (CCCP). Staining with MitoTracker Deep Red (MitoTracker DR, a dye used to measure mitochondrial mass that is sensitive to mitochondrial membrane potential) revealed lower mitochondrial mass/membrane potential in AMPKα1/α2-deficient Treg cells relative to controls at homeostasis (**Fig 5D**) and in the lung during influenza pneumonia (**Fig 5E**), consistent with their impaired maximal OCR at homeostasis. AMPK also promotes glycolysis (*36*); however, the baseline and maximal extracellular acidification rate (ECAR, a measure of glycolytic rate) of AMPKα1/α2-deficient and control Treg cells was comparable between genotypes, suggesting that AMPK is not required to sustain glycolysis in Treg cells at homeostasis (**Fig 5F-G**). AMPK also promotes autophagy through inhibition of mammalian target of rapamycin complex 1 (mTORC1) (*37, 38*), yet we found by flow cytometry that AMPKα1/α2- deficient Treg cells had no significant differences in expression of the autophagy marker LC3B (**Fig 5H**). These results collectively suggest that Treg cells require AMPK to maximize their mitochondrial mass and electron transport chain function.

**Figure 5.**
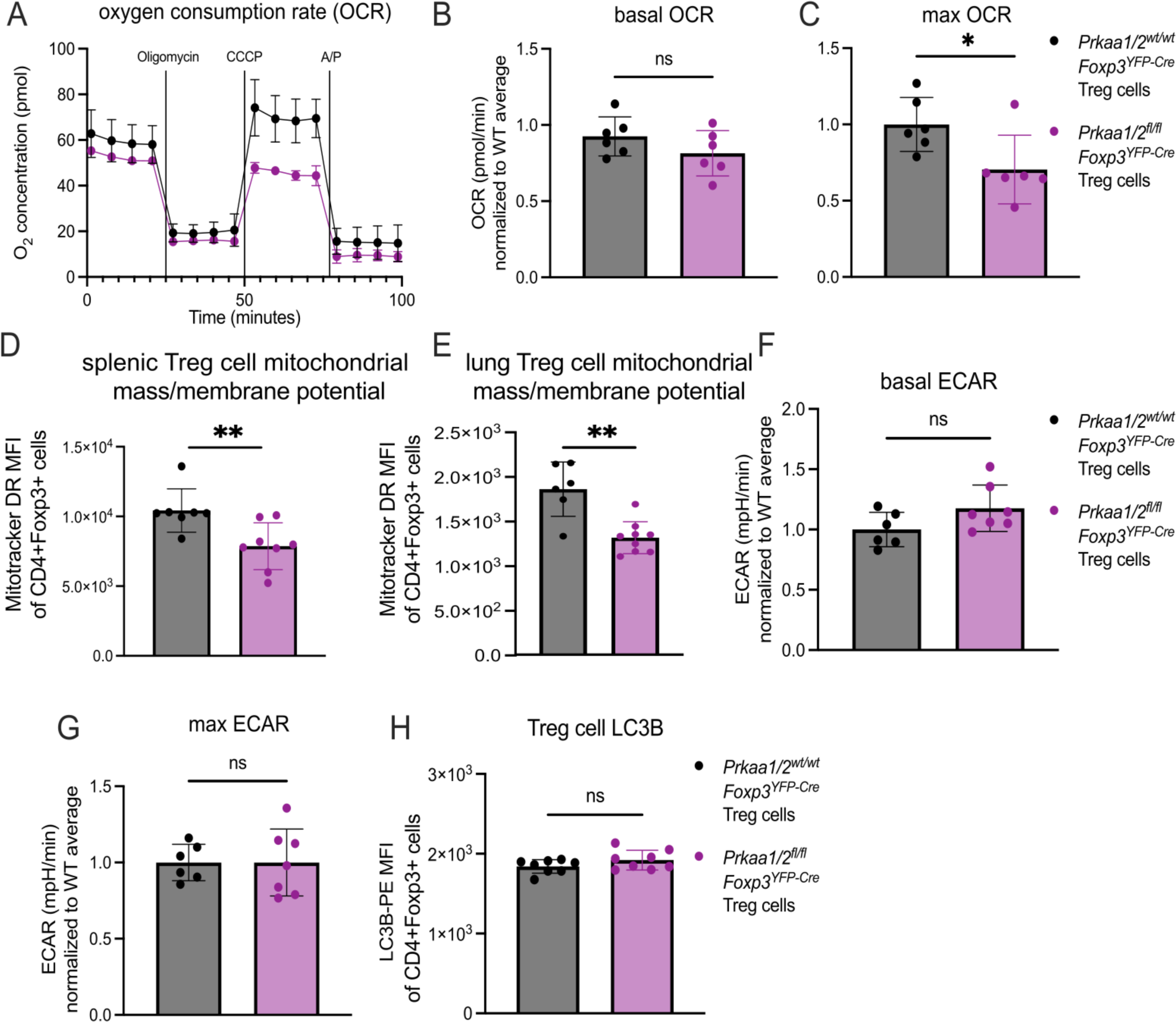
Treg cell AMPKα is necessary for maximal mitochondrial function. (**A**) Representative oxygen consumption rate (OCR) over time of CD4+Foxp3+ splenocytes from *Prkaa1/2^wt/wt^Foxp3^YFP-Cre^*(control, *n*=3) and *Prkaa1/2^fl/fl^Foxp3^YFP-Cre^*(*n*=2) mice following treatment of oligomycin (2.5 uM), carbonyl cyanide m-chlorophenylhydrazone (CCCP; 10 uM), and Antimycin A/Piercidin (A/P; 2 uM each) as measured by a metabolic flux assay. (**B**-**C**) Basal (B) and maximal (C) OCR of CD4+Foxp3+ splenocytes from control (*n*=6) and *Prkaa1/2^fl/fl^Foxp3^YFP-Cre^* (*n*=6) mice, some of which are shown in (A). (**D**-**E**) MitoTracker Deep Red (MitoTracker DR) mean fluorescence intensity (MFI) of CD4+Foxp3+ splenocytes at homeostasis (D; *n*=7 control, *n*=8 *Prkaa1/2^fl/fl^Foxp3^YFP-Cre^* mice) and lung CD4+Foxp3+ cells at day 10 post-influenza virus inoculation (E; same cohort as in Fig 4E-H and Supplemental Fig 6C-K, M-R, *n*=6 control, *n*=9 *Prkaa1/2^fl/fl^Foxp3^YFP-Cre^* mice). (**F**-**G**) Basal (F) and maximal (G) extracellular acidification rate (ECAR) of CD4+Foxp3+ splenocytes from control (*n*=6) and *Prkaa1/2^fl/fl^Foxp3^YFP-Cre^* (*n*=7) mice. (**H**) LC3B-PE MFI of CD4+Foxp3+ splenocytes from control (*n*=8) and *Prkaa1/2^fl/fl^Foxp3^YFP-Cre^*(*n*=8) mice. * *p* < 0.05, ** *p* < 0.01, ns not significant according to Mann-Whitney *U* test (**B**-**I**). Summary plots show all data points with mean and SD.

### AMPKα regulates DNMT1 to promote demethylation of metabolic genes

In human umbilical vein endothelial cells and mesenchymal stem cells cultured *in vitro*, AMPK phosphorylates DNMT1 to promote transcription of metabolic genes, including *Ppargc1a* (*17, 18*). Hence, we hypothesized that the lower expression of metabolic genes by tumor-infiltrating AMPKα1/α2-deficient Treg cells (see Cluster 1 in **Fig 2C-E**) was a consequence of DNA hypermethylation at their gene promoters. We tested this hypothesis by performing genome-wide DNA methylation profiling of *Prkaa1/2^fl/fl^Foxp3^YFP-Cre^*and control Treg cells sorted from B16 melanoma tumors and from spleens at homeostasis. While there was no difference in genome-wide promoter methylation, we observed hypermethylation of Cluster 1 gene promoters in AMPKα1/α2-deficient tumor-infiltrating Treg cells, as well as hypermethylation of *Ppargc1a* (PGC-1α) in both tumor-infiltrating and splenic AMPKα1/α2-deficient Treg cells (**Fig 6A-C**). Interestingly, we found that AMPKα1/α2-deficient splenic Treg cells express more DNMT1 protein relative to controls with no differences in *Dnmt1* gene expression, suggesting that AMPKα is a negative regulator of DNMT1 protein levels in Treg cells (**Fig 6D-E**). Co-immunoprecipitation assays in primary mouse iTreg cells, Jurkat cells, and the Treg cell-like MT-2 cell line (*39*) identified a physical interaction between AMPKα1 and DNMT1 (**Fig 6F** and **Supplemental Fig 7A-B**). To determine whether AMPK is present in the nucleus where it can interact with DNMT1, we performed immunofluorescence imaging in iTreg cells as well as Jurkat and MT-2 cells. Consistent with their physical interaction, our imaging studies identified AMPKα1 in the nucleus (**Fig 6G** and **Supplemental Fig 7C-D**). The subcellular compartmentalization of AMPKα1 was unaffected by activation with 5-aminoimidazole-4-carboxamide ribonucleoside (AICAR) in MT-2 cells (**Supplemental Fig 7E**). Finally, we established the functional significance of these findings by demonstrating that inhibition of DNMT activity with decitabine (DAC)— a clinically-used agent we showed in published work promotes Treg cell function and is sufficient to induce DNA hypomethylation in Treg cells (*32*)—increased MitoTracker DR staining (mitochondrial mass/membrane potential) in AMPKα1/α2-sufficient splenic Treg cells in a dose-dependent manner (**Fig 6H**). Treatment of AMPKα1/α2-deficient Treg cells with DAC also rescued MitoTracker DR signal to that of untreated control Treg cells, confirming that DNA methylation regulates mitochondrial mass in AMPKα1/α2-deficient Treg cells. Altogether, these experimental data reveal AMPK as a nuclear factor that regulates DNMT1 protein level in Treg cells to promote expression of metabolic factors that potentiate mitochondrial metabolism.

**Figure 6.**
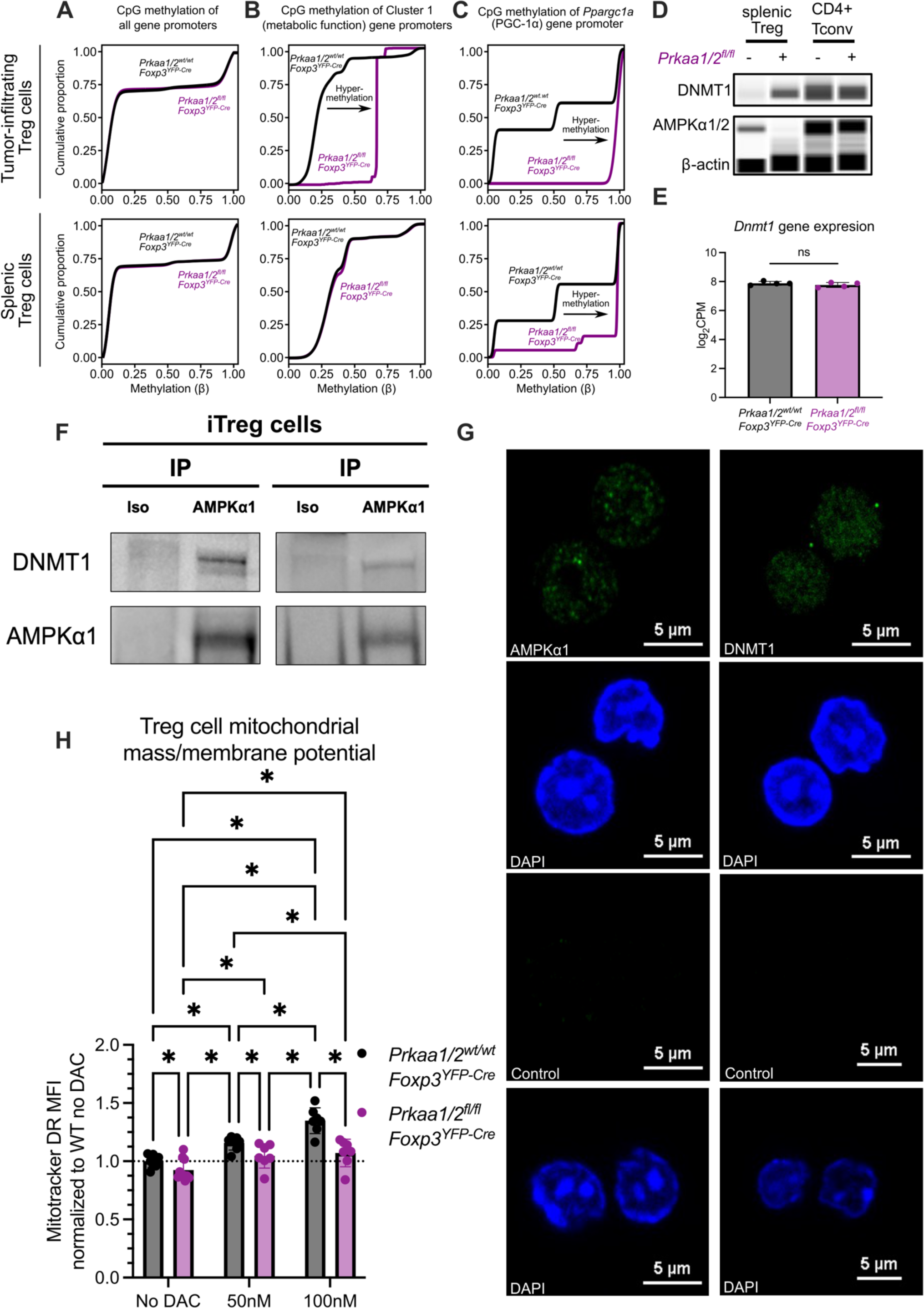
AMPKα1 interacts with DNMT1 to demethylate the promoter of mitochondrial genes in tumor-infiltrating Treg cells. (**A**-**C**) CpG methylation of all gene promoters (A), gene promoters of cluster 1 genes identified by *k*-means clustering of the RNA-sequencing shown in Fig 2C-F (B), and the *Ppargc1a* promoter (C) in tumor-infiltrating CD4+Foxp3+ cells (*n*=4 *Prkaa1/2^wt/wt^Foxp3^YFP-Cre^* or control, *n*=2 *Prkaa1/2^fl/fl^Foxp3^YFP-Cre^*) and splenic CD4+Foxp3+ cells at homeostasis (*n*=3 control, *n*=3 *Prkaa1/2^fl/fl^Foxp3^YFP-Cre^*) (**D**) DNMT1 protein expression of splenic CD4+Foxp3+ (Treg) and CD4+Foxp3-(CD4+ Tconv) cells at homeostasis (*n*=3 control, *n*=3 *Prkaa1/2^fl/fl^Foxp3^YFP-Cre^*; biological replicates were pooled and run as a single well). (**E**) *Dnmt1* gene expression of splenic CD4+Foxp3+ cells at homeostasis (*n*=4 control, *n*=4 *Prkaa1/2^fl/fl^Foxp3^YFP-Cre^*) as measured by RNA-sequencing shown in Fig 1. (**F**) anti-AMPKα1 and isotype control immunoprecipitates from *ex vivo* induced (i)Treg cell lysates blotted for DNMT1 protein. (**G**) Representative microscopy images of AMPKα-sufficient and-deficient iTreg cells showing AMPKα1 and DNMT1 subcellular localization. (**H**) Mitotracker Deep Red (Mitotracker DR) mean fluorescence intensity (MFI) of AMPKα-sufficient (control) and -deficient splenic CD4+Foxp3+ cells treated with either vehicle (*n*=8 control, *n*=10 *Prkaa1/2^fl/fl^Foxp3^YFP-Cre^*), 50 nM decitabine (DAC, *n*=7 control, *n*=7 *Prkaa1/2^fl/fl^Foxp3^YFP-Cre^*), or 100 nM DAC (*n*=7 control, *n*=7 *Prkaa1/2^fl/fl^Foxp3^YFP-Cre^*). * *p or q* < 0.05, ns not significant according to Mann-Whitney *U* test (**E**) with two-stage linear step-up procedure of Benjamini, Krieger, and Yekutieli with Q = 5% (**H**). Summary plots show all data points with mean and SD.

## Discussion

Treg cells exhibit metabolic plasticity in the TME, which in turn supports Treg cell suppressive function (*12, 13*). Nevertheless, mechanisms that orchestrate the metabolic adaptation of tumor-infiltrating Treg cells remain undetermined. Here, our experimental data revealed that AMPK-deficient Treg cells failed to exert optimal suppressive function in metabolically stressed microenvironments. We found that AMPK-deficient Treg cells were unable to augment their oxygen consumption under the stress of a mitochondrial uncoupling agent *ex vivo*, failed to upregulate genes supporting mitochondrial metabolism in the TME, and did not sustain proper mitochondrial mass/membrane potential or metabolic homeostasis during viral pneumonia. These results credential AMPK as a key mediator of Treg cell metabolic adaptation to settings of microenvironmental stress, likely through potentiation of mitochondrial metabolism, and are consistent with *in vitro* experiments suggesting AMPK potentiates Treg cell suppressive function (*40*).

While comparison of the metabolomic profiles of the TME and virus-injured lung interstitial fluid showed similar alterations when compared with plasma, we observed differences in the abundance of a small set of metabolites between these two interstitial fluid compartments, including glucose and glutamine, which were lower in the TME compared with the lung. In most cell types, increases in the AMP-to-ATP ratio from glucose deprivation and other states of energy stress lead to phosphorylation of AMPK by Liver kinase B1 (LKB1) (*41*). Notably, Treg cells require LKB1 to sustain immune self-tolerance at homeostasis, albeit in an AMPK signaling-independent manner (*19, 21*). Cell signaling events such as T cell receptor (TCR) engagement also activate AMPK via Calcium/Calmodulin dependent protein kinase kinase (CaMKK) (*16, 42*). These AMPK-activating events likely contribute to the metabolic adaptation mediated by Treg cell AMPK in disease microenvironments and serve as independent inputs through which AMPK can sense and respond to the extracellular milieu. Therefore, it is plausible that the loss-of-function we observed in tumor-infiltrating AMPK-deficient Treg cells is driven by an inability to adapt to glucose or other nutrient deprivation, whereas lung Treg cells require AMPK during influenza to adapt to different metabolic and signaling challenges. It nevertheless remains unclear what dimensions of Treg cell function are lost in AMPK-deficient Treg cells in these microenvironments, as our measurements of classical surface molecules via which Treg cells exert their suppressive function did not reveal broad changes in Treg cell suppressive phenotype.

Our experimental data suggest that AMPK regulates DNMT1 protein level to activate the expression of metabolic genes that support mitochondrial function, including *Ppargc1a*/PGC-1α. In some cell types cultured *in vitro*, an AMPK-DNMT1-mitochondrial metabolism axis regulates metabolic function (*17, 18*). *In vivo*, we found in tumor-infiltrating Treg cells that AMPK serves as an epigenetic regulator of transcriptional programs that support metabolic function. Loss of AMPK in Treg cells led to greater DNMT1 protein expression, which was associated with DNA hypermethylation at the promoters of key metabolic genes in the TME. Our co-immunoprecipitation studies confirmed that AMPKα1 directly interacts with DNMT1, likely regulating DNMT1 level and activity via phosphorylation events suggested by *in vitro* studies (*17, 18*). Critically, treatment with the DNMT inhibitor decitabine rescued mitochondrial mass in AMPK-deficient Treg cells, mechanistically connecting AMPK, DNMT1, and mitochondria. Additional mechanisms may link AMPK to DNA methylation writer complexes. For example, UHRF1, the DNMT1 adapter protein we previously showed to be necessary for Treg cell identity and function (*23*), was recently reported to inhibit AMPK function in the nucleus of hepatocytes (*43*). Hence, AMPK may regulate DNA methylation in Treg cells via interaction with other DNMT complex members such as UHRF1.

The context-specific upregulation of *Prkaa2* in tumor-infiltrating Treg cells may explain the discrepant consequences for anti-tumor immunity reported in Treg cell-specific AMPKα1-deficient mice challenged with B16 melanoma tumors (*27, 28*). A study assessing the contribution of each AMPK catalytic subunit isoform to the potentiation of mitochondrial gene expression found that AMPKα2, but not AMPKα1, is required for the upregulation of *Ppargc1a* expression during myotube differentiation (*44*). While we showed that AMPKα1 also interacts with DNMT1 in T cell lines and primary FOXP3^+^ T cells, it is plausible that the two AMPKα isoforms exert differential regulation over epigenetic modifiers in the TME. Therefore, *Prkaa2* upregulation by tumor-infiltrating, AMPKα1-deficient Treg cells may impact Treg cell suppressive function and thereby lead to conflicting results, especially if *Prkaa2* upregulation is modified by variables that are difficult to control across studies, such as the mouse colony microbiome (*45*). Indeed, our data suggest that, while AMPKα1 and AMPKα2 may have a shared set of downstream targets that are necessary for Treg cell function in the TME, isoform-specific activities may have divergent influences on Treg cell function.

A recent clinical trial showed that metformin, an indirect AMPK activator, significantly reduces the risk of developing long COVID (*46-48*). This finding is consistent with the observed effects of metformin on mouse models of lung injury (*49, 50*). Our data support that AMPK is dispensable for Treg cell-mediated immune self-tolerance yet promotes Treg cell suppressive function in disease microenvironments. This context-specific requirement of AMPK for Treg cell function makes it an attractive drug target for attempts to potentiate the function of Treg cells *ex vivo* before their use in cell-based therapies, such as those being leveraged in early phase clinical trials to improve outcomes in patients with COVID-19 (*51, 52*).

Our study has limitations. First, AMPK phosphorylates specific residues of DNMT1 in human umbilical vein endothelial cells to decrease DNMT1 function (*17*). Unfortunately, antibodies specific for the homologous residues of mouse DNMT1 are not available. Regardless, our co-immunoprecipitation, immunofluorescence, immunoassay, and sequencing data support that AMPK regulates DNMT1 level and function in Treg cells. Second, we detected 159 metabolites via LC-MS in ∼5×10^4^ Treg cells sorted from the influenza virus-injured lung at peak injury. While we were able to detect an accumulation of pyruvate and lactate in AMPK-deficient Treg cells suggestive of an impaired TCA cycle, a more comprehensive assessment of the Treg cell metabolome during viral pneumonia may have provided insight into whether the loss-of-function in this context is due to energy stress in the absence of AMPK-mediated metabolic adaptation. Finally, the loss of AMPK-dependent regulation of transcriptomic and epigenetic signatures may be too complex to cause the resulting Treg cell loss-of-function via a single factor, such as dampened *Ppargc1a* expression; the combined dysregulation of more than one downstream target of AMPK is likely to mediate the loss of function.

In summary, our findings support a model in which AMPK coordinates the metabolic adaptation of Treg cells in settings of microenvironmental stress by potentiating mitochondrial metabolism, consistent with AMPK’s canonical function as a sensor of energetic stress and the central role mitochondrial metabolism plays in programming Treg cell functional state. We show that this AMPK-mediated metabolic adaptation is executed in part through the regulation of DNA methylation at key metabolic loci, offering potential pharmacologic targets to modulate Treg cell function in disease, including in severe lung injury and cancer.

## Methods

### Mice

All mouse procedures were approved by the Northwestern University IACUC under protocols IS00012519 and IS00017837. *Prkaa1^fl/fl^*(cat. no. 014141)*, Prkaa2^fl/fl^* (cat. no. 014142), and *Foxp3^YFP-Cre^* (cat. no. 016959) mice from the C57BL/6J genetic background were purchased from The Jackson Laboratory and bred to generate Treg cell-specific AMPKα1 and AMPKα2 double KO mice (*Prkaa1^fl/fl^xPrkaa2^fl/fl^xFoxp3^YFP-Cre^),* and Treg cell-specific AMPKα1 and AMPKα2 single KO mice (*Prkaa1^fl/fl^xFoxp3^YFP-Cre^*and *Prkaa2^fl/fl^xFoxp3^YFP-Cre^*, respectively). All animals were genotyped using services provided by Transnetyx Inc., with primers provided by The Jackson Laboratory and shown in **Supplemental Table 1**; RT-PCR primers are also shown here. Animals received water *ad libitum*, were housed at a temperature range of 20°C–23°C under 14-hour light/10-hour dark cycles and received standard rodent chow.

### Flow cytometry and cell sorting

Single-cell suspensions of organ tissues, blood, tumors, or cultured cells were prepared and stained for flow cytometry analysis and sorting as previously described (*23, 33*) using the reagents shown in **Supplemental Table 2**. Cell counts of single-cell suspensions were obtained using a Cellometer with AO/PI staining ((Nexcelom Bioscience cat. # SD014-0106)) before preparation for flow cytometry. Data acquisition for analysis was performed using a BD LSRFortessa or Symphony A5 instrument with FACSDiva software (BD). Cell sorting was performed using the 4-way purity setting on BD FACSAria SORP instruments with FACSDiva software or with a microfluidics MACSQuant Tyto sorter (Miltenyi). Analysis was performed with FlowJo, version 10.9.0 software. Dead cells were excluded using a viability dye for analysis and sorting (*53*).

### iTreg cell induction and culture

CD4+CD25- T cells were enriched from splenocyte single cell suspensions using the negative fractions from the Miltenyi mouse CD4+CD25+ Regulatory T Cell Isolation Kit (cat. no. 130-091-041) according to manufacturer instructions or sorted via flow cytometry. CD4+CD25- T cells were seeded in a 24-well plate at a concentration of 3×10^5^ cells/mL and cultured for 3-5 days in complete RPMI medium (10% FBS) containing recombinant IL-2 (50IU/mL) and TGF-β (10 ng/mL) (*23*). *Foxp3*-YFP+ cells were sorted using flow cytometry for downstream use.

### B16 melanoma tumor model

B16-F10 cells (ATCC CRL-6475) were cultured in RPMI (10% FBS, 5% L-glutamine, 5% sodium pyruvate, 5% penicillin/streptomycin, 2.5% HEPES buffer), harvested at 60- 70% confluence, and counted using a Cellometer with AO/PI staining (Nexcelom Bioscience cat. # SD014-0106) as previously described (*23*). 250,000 B16-F10 cells were resuspended in 0.1 mL of PBS and 40% Matrigel (Corning cat. # 356237) and injected subcutaneously in the hair-trimmed flanks of 12–15-week-old mice. Tumor length and width were measured every other day starting day 7 post-engraftment up to day 21 post-engraftment. A subset of tumors was resected post-mortem for flow cytometry analysis and sorting as follows: subcutaneous tumors were harvested from euthanized mice and minced with surgical scissors in 3mL of HBSS containing 2 mg of collagenase (Sigma Aldrich cat. no. 11088866001) and 0.25 mg of DNAse I (Sigma Aldrich cat. no. 10104159001) per mL. Tumor homogenates were incubated at 37°C for 45 min, transferred to gentleMACS™ C Tubes (Miltenyi Biotec cat. # 130-093-237) and processed with the mouse implanted tumor protocol (m_impTumor_01). Tumor homogenates were then subjected to debris removal (Miltenyi cat. # 130-109-398) and dead cell removal (Miltenyi cat. # 130-090-101) according to the kit manufacturer’s instructions. The debris and dead cell-depleted tumor homogenates were then stained for flow cytometry and cell sorting.

### Influenza A virus administration

Mice were anesthetized with isoflurane and intubated using a 20- gauge angiocatheter cut to a length that placed the tip of the catheter above the carina. Mice were instilled with mouse-adapted influenza A/WSN/33 [H1N1] virus (12.5 pfu in 50 μL of sterile PBS) as previously described (*33*).

### Measurement of physiologic readouts of influenza pneumonia progression and resolution

Arterial blood oxygen saturation (SpO2) was measured in control and influenza virus-infected mice using a MouseOx Plus pulse oximeter (Starr Life Sciences). Beginning on the fifth day post-infection and continuing every other, SpO2 was measured with oximeter collar clips secured to the hairless neck of conscious, immobilized animals. Mouse weights were recorded the day of influenza virus inoculation and every other day post-inoculation starting day 5. Mouse weights were normalized to those recorded on the day of inoculation.

### Lung tissue harvesting and processing

Influenza virus-infected mice were euthanized and slowly infused with HBSS through the right atrium of the heart, clearing the pulmonary circulation of blood. The lungs were harvested and grossly homogenized with scissors in HBSS containing 2 mg of collagenase D (Sigma Aldrich cat. no. 11088866001) and 0.25 mg of DNAse I (Sigma Aldrich cat. no. 10104159001) per mL, incubated for 45min at RT, and then further homogenized using the mouse lung protocol of the Miltenyi OctoMACS tissue dissociator (m_lung_02). These procedures have been previously reported (*33*).

### Western blotting

Cultured cells were lysed for one hour at 4°C in lysis buffer (Cell Signaling cat. no. 9803) supplemented with phosphatase (Cell Signaling cat. no. 5870S) and protease inhibitors (Roche, cat. no. 65726900) after which their concentration was measured with a BCA assay according to manufacturer instructions (Pierce cat. no. 23225). Cell lysates were subjected to gel electrophoresis and transferred to membranes that were incubated with an antibody against AMPKα1 (Abcam cat. no. ab32047), DNMT1 (Cell Signaling cat. no. 5032) and β-actin (Abcam cat. no. ab8227) overnight at 4 °C with constant agitation.

### Wes protein immunoassay

Flow cytometry sorted cells were lysed and the resulting lysate protein concentrations were measured as described above. For protein measurements using the Simple Wes immunoassay system, 0.5 ug of protein in 3 uL were loaded per well and processed according to manufacturing instructions. The following concentrations were used for primary antibodies: 1:50 anti-DNMT1 (Invitrogen cat. no. MA5-16169), 1:50 anti-AMPKα (Cell Signaling cat. no. 2532S), and 1:50 anti-β-actin (Abcam cat. no. ab8227).

### Co-immunoprecipitation assay

10^6^ cells were lysed in cell lysis buffer for one hour at 4°C as described above. Lysates were incubated with an antibody against AMPKα1 (Abcam cat. no. ab32047) or isotype control (Cell Signaling cat. no. 7074) overnight at 4 °C with constant agitation. The immune complex was precipitated with Dyna Protein G beads (Life Technologies cat. no. 10003D), washed and resuspended in SDS/PAGE loading buffer, and heated to 95 °C for 5 minutes. Processed samples were then blotted with antibodies against DNMT1 (Cell Signaling cat. no. 5032), AMPKα1 (Abcam cat. no. ab32047), and β-actin (Abcam cat. no. ab8227).

### Immunofluorescence for microscopy

10^6^ cells were fixed with ice-cold 100% methanol for 5 minutes. Subsequently, samples were processed with Immunofluorescence Application Solutions Kit (Cell Signaling cat. no. 12727) following the manufacturer’s protocol. Cells were stained overnight at 4°C with anti-DNMT1 (Abcam cat. no. ab21799 1), anti-AMPKα1 (Abcam cat. no. ab32047), Alexa fluor 488-conjugated isotype control (Abcam cat. no. ab199091), or unconjugated isotype control (Abcam cat. no. ab172730). The following day, cells that were stained with anti-AMPKα1 antibody and unconjugated isotype control antibody were incubated in the dark at room temperature for 2 hours with anti-rabbit Alexa Fluor 488 secondary antibody (Abcam cat. no. ab150113). Following antibody incubation, cells were mounted on a slide with VECTASHIELD Vibrance mounting medium containing DAPI (Vector Labs cat. no. H-1800). Fluorescent images were acquired at room temperature using a confocal microscope (Nikon) with 40× magnification at the Northwestern Center for Advanced Microscopy.

### Nuclear-cytoplasmic fractionation assay

5 x 10^6^ MT-2 cells were treated and subsequently underwent lysis using NE-PER™ Nuclear and Cytoplasmic Extraction kit (ThermoFisher cat. no. 78833) according to manufacturer’s protocol. Nuclear and cytoplasmic fractions were collected and further analyzed for the expression of proteins of interest with Western Blotting as described above.

### Metabolic flux (seahorse) assay

2.5 x 10^5^ flow cytometry-sorted Treg cells were seeded on a 96- well Seahorse cell culture plate and analyzed on a Seahorse XF24 Analyzer (*10*). The following drugs and corresponding doses were loaded onto ports A, B, C, and D in the same order: Oligomycin (2.5 uM, Sigma-Aldrich cat no. 75351), CCCP (10 uM, Sigma-Aldrich cat no. C2759), Antimycin A/Piercidin (2 uM each, Sigma-Aldrich cat no. A8674 and 15379, respectively), and 2-deoxyglucose (25 mM, Sigma-Aldrich cat no. D8375).

### RNA-sequencing, modified reduced representation bisulfite sequencing (mRRBS) and analysis

10^3^-10^5^ flow cytometry sorted cells were lysed immediately after sorting with QIAGEN RLT Plus containing 1% β-mercaptoethanol and subjected to RNA and DNA isolation using the QIAGEN AllPrep Micro Kit as previously described. RNA-seq library preparation was performed using the SMARTer Stranded Total RNA-Seq Kit, version 2 (Takara cat. no. 634411) and mRRBS library preparation was performed using custom procedures previously described by our group (*23, 33, 54*). After sequencing, raw binary base call (BCL) files were converted to FASTQ files using bcl2fastq (Illumina). All FASTQ files were processed using the nf-core/RNA-seq pipeline (*55*) version 3.9 implemented in Nextflow (*56, 57*) 23.04.2 with Northwestern University Quest HPC (Genomic Nodes) configuration (nextflow run nf-core/rnaseq-profile nu_genomics --genome GRCm38). In short, lane-level reads were trimmed using trimGalore! 0.6.5, aligned to the GRCm38 reference genome using STAR 2.7.11, and quantified using Salmon. All samples showed satisfactory alignment rate (>60%). After quantification, differential expression analysis was performed in R version 4.2.0 using DEseq2 v1.38.3 (*58*). Sample genotype was used as an explanatory factor. Within-group homogeneity was first confirmed by principal component analysis (PCA) and no outliers were found. K-means clustering of differentially expressed genes (*q* < 0.05) was performed using a previously published custom R function (*59*). In brief, k was first determined using the elbow plot and the kmeans function in R stats 3.6.2 (Hartigan–Wong method with 25 random sets and a maximum of 1,000 iterations) was used for k-means clustering. Samples were clustered using Ward’s method and a heatmap was generated using pheatmap version 1.0.12. Subsequent GO term enrichment was performed using topGO version 2.50.0 with Fisher’s exact test. org.Mm.eg.db version 3.16.0 and GO.db version 3.16.0 was included in GO terms enrichment analysis as references. Gene Set Enrichment Analysis was performed using the GSEA, version 4.0.3, GSEAPreranked tool with genes ordered by log2(fold change) in average expression (*60*). mRRBS analysis was performed using previously published procedures (*61*).

### Collection of lung Treg cells for metabolomics

Lungs were harvested and processed as described above. Lung single cell suspensions were subjected to CD4+ cell positive enrichment according to kit manufacturer’s instructions (Miltenyi Biotec, cat. no. 130-097-048) before fluorochrome staining. Fluorochrome stained cells were then resuspended in sorting buffer (Miltenyi Biotec, cat. no. 130-107- 207) to achieve a concentration of 2 x 10^6^ cells / mL of buffer. Using a MACSQuant Tyto, 5-10 x 10^5^ lung Treg cells were sorted from each pair of lungs. Sorted cells were centrifuged at 500 rcf for 6 minutes, 4 °C. After the sorting media was removed via pipetting, the pelleted cells were resuspended in 15 µl of 80% acetonitrile and vortexed for 30 seconds. Following centrifugation for 30 min at 20,000 f, 4 °C, the supernatant was collected for High-Performance liquid chromatography and high-resolution mass spectrometry and tandem mass spectrometry (LC-MS) analysis.

### Collection of interstitial fluid and plasma for metabolomics

Blood was collected from both influenza-infected and B16-melanoma tumor-engrafted mice via insertion of glass capillaries through the retro-orbital sinus. This blood was centrifuged at 800 rcf for 10 minutes, 4 °C in EDTA tubes. The plasma phase was pipetted, frozen with liquid nitrogen, and stored at −80 °C. Subcutaneous B16-F10 melanoma tumors were harvested on day 15 post-engraftment. Lungs of mice that received 12.5 PFUs of influenza A virus intra-tracheally were harvested on day 10 post-inoculation. The tumors were rinsed in phosphate buffered saline and dried with task wipers. The lungs were perfused via injection of 10 mL of Hanks Balanced Salt Solution (HBSS) through the heart’s right ventricle before harvesting all soft organs of the mediastinum. The mediastinal structures surrounding the lungs were removed via gross dissection. The intact tumors and lungs were then centrifuged at 100 rcf for 10 minutes, 4 °C in centrifuge tubes containing a 0.22 µm filter (Costar cat. no. 8160). The extracted interstitial fluid was then diluted 1 to 5 in 80% acetonitrile and vortexed for 30 seconds. The diluted interstitial fluid was centrifuged for 30 min at 20,000 rcf, 4 °C and the supernatant was collected for LC-MS analysis.

### High-Performance liquid chromatography and high-resolution mass spectrometry and tandem mass spectrometry (LC-MS) for metabolomics

Samples were analyzed by LC-MS. Specifically, system consisted of a Thermo Q-Exactive in line with an electrospray source and an Ultimate3000 (Thermo) series HPLC consisting of a binary pump, degasser, and auto-sampler outfitted with a Xbridge Amide column (Waters; dimensions of 3.0 mm × 100 mm and a 3.5 µm particle size). The mobile phase A contained 95% (vol/vol) water, 5% (vol/vol) acetonitrile, 10 mM ammonium hydroxide, 10 mM ammonium acetate, pH = 9.0; B was 100% Acetonitrile. The gradient was as following: 0 min, 15% A; 2.5 min, 30% A; 7 min, 43% A; 16 min, 62% A; 16.1-18 min, 75% A; 18-25 min, 15% A with a flow rate of 150 μL/min. The capillary of the ESI source was set to 275 °C, with sheath gas at 35 arbitrary units, auxiliary gas at 5 arbitrary units and the spray voltage at 4.0 kV. In positive/negative polarity switching mode, an *m*/*z* scan range from 60 to 900 was chosen and MS1 data was collected at a resolution of 70,000. The automatic gain control (AGC) target was set at 1 × 10^6^ and the maximum injection time was 200 ms. The top 5 precursor ions were subsequently fragmented, in a data-dependent manner, using the higher energy collisional dissociation (HCD) cell set to 30% normalized collision energy in MS2 at a resolution power of 17,500. Besides matching m/z, metabolites are identified by matching either retention time with analytical standards and/or MS2 fragmentation pattern. Data acquisition and analysis were carried out by Xcalibur 4.1 software and Tracefinder 4.1 software, respectively (both from Thermo Fisher Scientific).

### LC-MS data analysis

Raw peak intensity data of the metabolites detected by LC-MS were uploaded to Metaboanalyst 5.0’s statistical analysis [one factor] module. For comparisons with more than two groups (**Fig 3**), one-way ANOVA with *q* < 0.05 was employed to identify significant differentially enriched metabolites. Comparisons with only two groups (**Fig 4, Supplemental Fig 4-5**) were analyzed with multiple parametric t-tests and fold-change analysis using Metabonalyst 5.0’s standard settings (*p* < 0.1). Fold change threshold was set to 1.5 in resulting volcano plots to increase the power of the downstream overrepresentation analysis. The names of significant differentially enriched metabolites were used as inputs for Metaboanalyst 5.0’s enrichment analysis module. The KEGG human metabolic pathway metabolite set library was selected for enrichment analysis.

### Statistical analysis

*p*-values and FDR *q*-values resulting from two-tailed tests were calculated using statistical tests stated in the figure legends using GraphPad Prism v10.1.0. Differences between groups with *p* or *q* values < 0.05 were considered statistically significant; see **LC-MS data analysis** for the statistical approach to metabolomic profiling data and **RNA-sequencing, modified reduced representation bisulfite sequencing (mRRBS) and analysis** for the statistical approach to transcriptomic and epigenomic profiling data. Central tendency and error are displayed as mean ± standard deviation (SD) except as noted. Box plots show median and quartiles. Numbers of biological replicates are stated in the figures or accompanying legends. Computational analysis was performed using Genomics Nodes and Analytics Nodes on Quest, Northwestern University’s High-Performance Computing Cluster.

### Data availability

Raw sequencing data will be made publicly available on the GEO database pending peer-reviewed publication.

## Acknowledgements.

MATA is supported by NIH awards T32GM144295, T32HL076139, and F31HL162490. NM is supported by NIH award T32AI083216. CPRF is supported by T32HL076139. QL is supported by the David W. Cugell Fellowship and the The Genomics Network (GeNe) Pilot Project Funding. AMJ is supported by NIH award F32HL162418. LM-N is supported by the NIH awards K08HL159356, U19AI135964 and the Parker B. Francis Opportunity Award. KC is supported by NIH award T32GM144295. SEW is supported by the Burroughs Wellcome Fund Career Awards for Medical Scientists. BDS is supported by NIH awards R01HL149883, R01HL153122, P01HL154998, P01AG049665, and U19AI135964. We wish to acknowledge the Northwestern University Flow Cytometry Core Facility supported by CA060553; the BD FACSAria SORP system was purchased with the support of S10OD011996. We also wish to acknowledge the Northwestern University RNA-Seq Center/Genomics Lab of the Pulmonary and Critical Care Medicine and Rheumatology Divisions. Histology services were provided by the Northwestern University Mouse Histology and Phenotyping Laboratory, which is supported by P30CA060553 awarded to the Robert H. Lurie Comprehensive Cancer Center. This research was supported in part through the computational resources and staff contributions provided by the Genomics Compute Cluster, which is jointly supported by the Feinberg School of Medicine, the Center for Genetic Medicine, and Feinberg’s Department of Biochemistry and Molecular Genetics, the Office of the Provost, the Office for Research, and Northwestern Information Technology. The Genomics Compute Cluster is part of Quest, Northwestern University’s high performance computing facility, with the purpose to advance research in genomics.

---

**Supplemental Figure 1.**
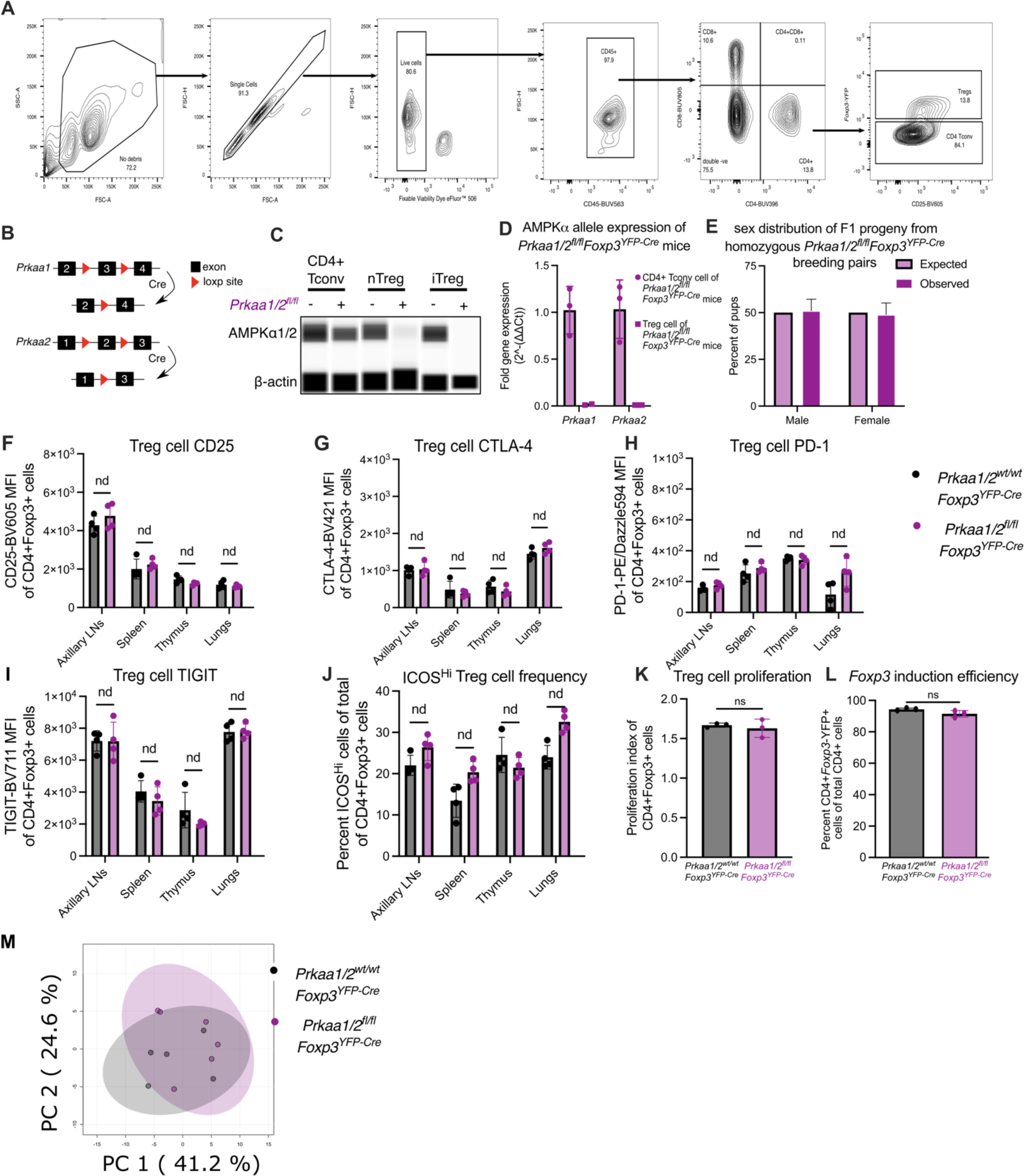
Validation of AMPKα1/α2 conditional knockout and Treg cell and mouse phenotyping of *Prkaa1/2^fl/fl^Foxp3^YFP-Cre^* and control mice. (**A**) Flow cytometry gating strategy for the phenotyping of T cell populations, including Treg cells (CD4+Foxp3+). (**B**) Schematic of the *Foxp3*- YFP-Cre-mediated excision of loxP-flanked exons of *Prkaa1* and *Prkaa2* in Foxp3+ cells. (**C**) Validation of Treg cell-specific KO of AMPKα1 and AMPKα2 using the Simple Wes protein immunoassay system (*n*=3 replicates per group and per cell type were pooled and run as a single well). (**D**) Relative expression of *Prkaa1* and *Prkaa2* by CD4+Foxp3-conventional T (CD4+ Tconv) cells and CD4+Foxp3+ Treg cells of *Prkaa1/2^fl/fl^Foxp3^YFP-Cre^*mice (*n*=3 CD4+Foxp3-cells, *n*=2 CD4+Foxp3+ cells). (**E**) Observed and expected frequencies of F1 pups resulting from crosses of male *Prkaa1/2^fl/fl^Foxp3^YFP-Cre/Y^* mice with female *Prkaa1/2^fl/fl^Foxp3^YFP-Cre/YFP-Cre^*mice. Chi-square test for goodness of fit p = 0.75. Chi-square = 0.1 with 1 degree of freedom (n=127 males and 122 females). (**F**-**I**) CD25-BV605 (F), CTLA-4-BV421 (G), PD-1-PE/Dazzle594 (H), and TIGIT-BV711 (I) mean fluorescence intensity (MFI) of CD4+Foxp3+ cells from *Prkaa1/2^wt/wt^Foxp3^YFP-Cre^*(control) and *Prkaa1/2^fl/fl^Foxp3^YFP-Cre^* mouse axillary lymph nodes (LNs), spleen, thymus, and lung (*n*=4 control, *n*=4 *Prkaa1/2^fl/fl^Foxp3^YFP-Cre^*). (**J**) Frequency of ICOS^Hi^ CD4+Foxp3+ cells of total CD4+Foxp3+ cells from control and *Prkaa1/2^fl/fl^Foxp3^YFP-Cre^* mouse LNs, spleen, thymus, and lung (*n*=4 control, *n*=4 *Prkaa1/2^fl/fl^Foxp3^YFP-Cre^*). (**K**) Proliferation index of Cell Trace Violet+CD4+Foxp3+ cells according to FlowJo 10.9.0’s proliferation modeling tool (*n*=3 control, *n*=3 *Prkaa1/2^fl/fl^Foxp3^YFP-Cre^*). (**L**) Frequency of CD4+Foxp3+ cells of total CD4+ cells after *in vitro* treatment of CD4+Foxp3-splenocytes with Treg cell-polarizing conditions for 5 days (*n*=3 control, *n*=3 *Prkaa1/2^fl/fl^Foxp3^YFP-Cre^*). (**M**) Principal component (PC) analysis of the LC-MS data generated with metabolites extracted from splenic Treg cells of control (*n*=5) and *Prkaa1/2^fl/fl^Foxp3^YFP-Cre^* (*n*=6) mice. Ns not significant, nd no discovery according to Mann-Whitney *U* test (**K**-**L**) with two-stage linear step-up procedure of Benjamini, Krieger, and Yekutieli with Q = 5% (**F**- **J**). Summary plots show all data points with mean and SD.

**Supplemental Figure 2.**
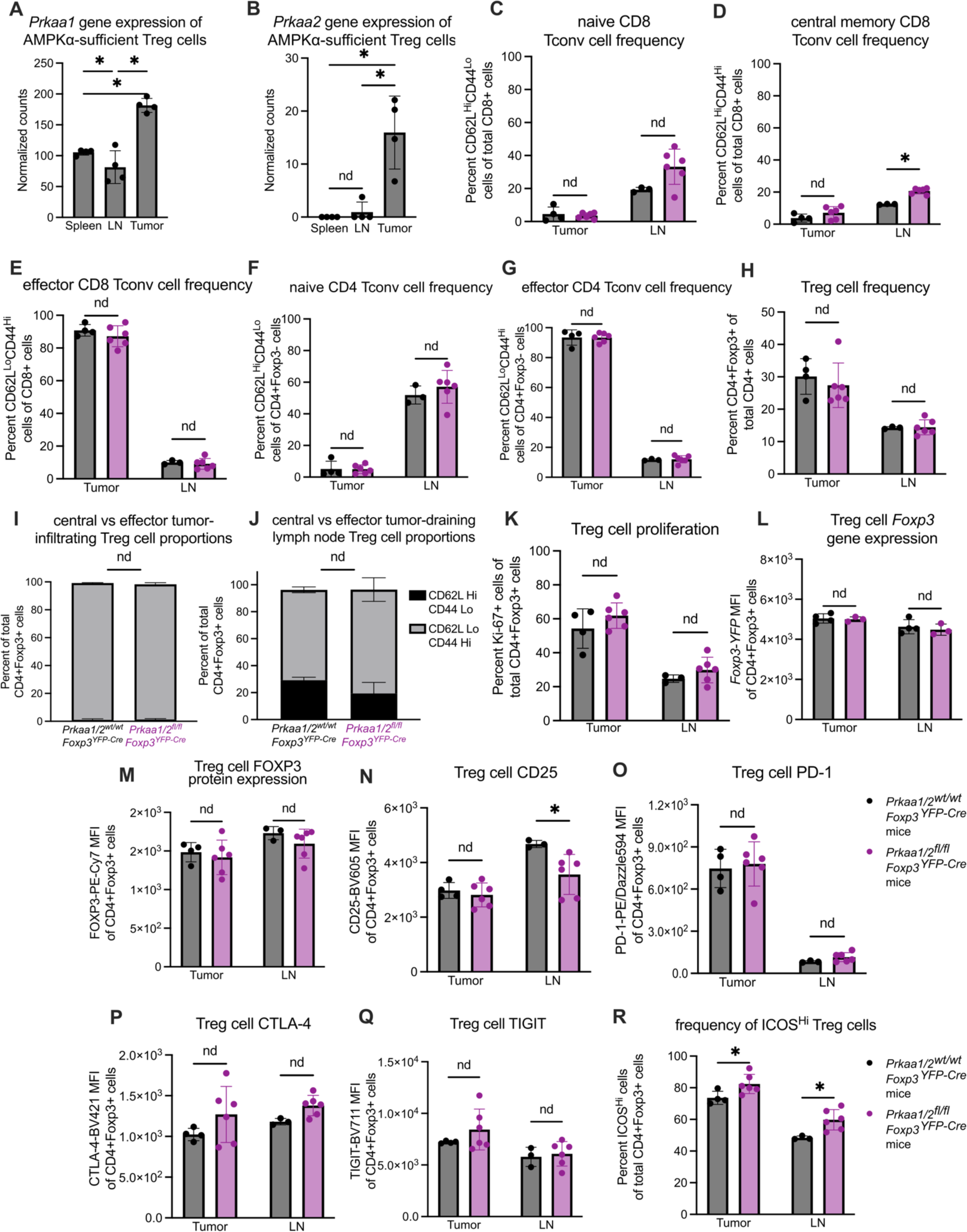
Immune phenotyping of tumor-infiltrating T cell subsets of *Prkaa1/2^fl/fl^Foxp3^YFP-Cre^*and control mice. (**A**-**B**) Normalized gene expression counts of *Prkaa1* (A) and *Prkaa2* (B) by CD4+Foxp3+ cells sorted from tumors, spleens, and lymph nodes of tumor-bearing *Prkaa1/2^wt/wt^Foxp3^YFP-Cre^*(control, *n*=4) mice. (**C**-**E**) Frequency of naive (CD62L^Hi^CD44^Lo^; C), central memory (CD62L^Hi^CD44^Hi^; D), and effector (CD62L^Lo^CD44^Hi^; E) CD8+ conventional T (Tconv) cells out of total CD8+ cells in tumors (*n*=4 control, *n*=6 *Prkaa1/2^fl/fl^Foxp3^YFP-Cre^*) and tumor-draining lymph nodes (LN; *n*=3 control, *n*=6 *Prkaa1/2^fl/fl^Foxp3^YFP-Cre^*mice). (**F**-**G**) Frequency of naive (F) and effector (G) CD4+Foxp3-Tconv cells out of total CD4+Foxp3-cells in tumors (*n*=4 control, *n*=6 *Prkaa1/2^fl/fl^Foxp3^YFP-Cre^* mice) and LN (*n*=3 control, *n*=6 *Prkaa1/2^fl/fl^Foxp3^YFP-Cre^* mice). (**H**) Percent of CD4+Foxp3+ cells out of total CD4+ cells in tumors (*n*=4 control, *n*=6 *Prkaa1/2^fl/fl^Foxp3^YFP-Cre^* mice) and LN (*n*=3 control, *n*=6 *Prkaa1/2^fl/fl^Foxp3^YFP-Cre^*mice). (**I**-**J**) Frequency of central (CD62L^Hi^CD44^Lo^) and effector (CD62L^Lo^CD44^Hi^) CD4+Foxp3+ cells out of total CD4+Foxp3+ cells in tumors (I; *n*=4 control, *n*=6 *Prkaa1/2^fl/fl^Foxp3^YFP-Cre^*mice) and LN (J; *n*=3 control, *n*=6 *Prkaa1/2^fl/fl^Foxp3^YFP-Cre^*mice). (**K**) Ki- 67+CD4+Foxp3+ cells out of total CD4+Foxp3+ cells in tumors (*n*=4 control, *n*=6 *Prkaa1/2^fl/fl^Foxp3^YFP-Cre^* mice) and LN (*n*=3 control, *n*=6 *Prkaa1/2^fl/fl^Foxp3^YFP-Cre^* mice). (**L**) *Foxp3*-YFP mean fluorescence intensity (MFI) of CD4+Foxp3+ cells in tumors (*n*=4 control, *n*=6 *Prkaa1/2^fl/fl^Foxp3^YFP-Cre^* mice) and LN (*n*=3 control, *n*=6 *Prkaa1/2^fl/fl^Foxp3^YFP-Cre^* mice). (**M**-**Q**) FOXP3-PE-Cy7 (M), CD25-BV605 (N), PD-1- PE/Dazzle594 (O), CTLA-4-BV421 (P) and TIGIT-BV711 (Q) MFI of CD4+Foxp3+ cells from tumors (*n*=4 control, *n*=6 *Prkaa1/2^fl/fl^Foxp3^YFP-Cre^* mice) and LN (*n*=3 control, *n*=6 *Prkaa1/2^fl/fl^Foxp3^YFP-Cre^* mice). (**R**) Frequency of ICOS^Hi^ CD4+Foxp3+ cells out of total CD4+Foxp3+ cells in tumors (*n*=4 control, *n*=6 *Prkaa1/2^fl/fl^Foxp3^YFP-Cre^* mice) and LN (*n*=3 control, *n*=6 *Prkaa1/2^fl/fl^Foxp3^YFP-Cre^* mice). * *q* < 0.05, nd no discovery according to Mann Whitney U test with two-stage linear step-up procedure of Benjamini, Krieger, and Yekutieli with Q = 5% (**A**-**R**). Summary plots show all data points with mean and SD.

**Supplemental Figure 3.**
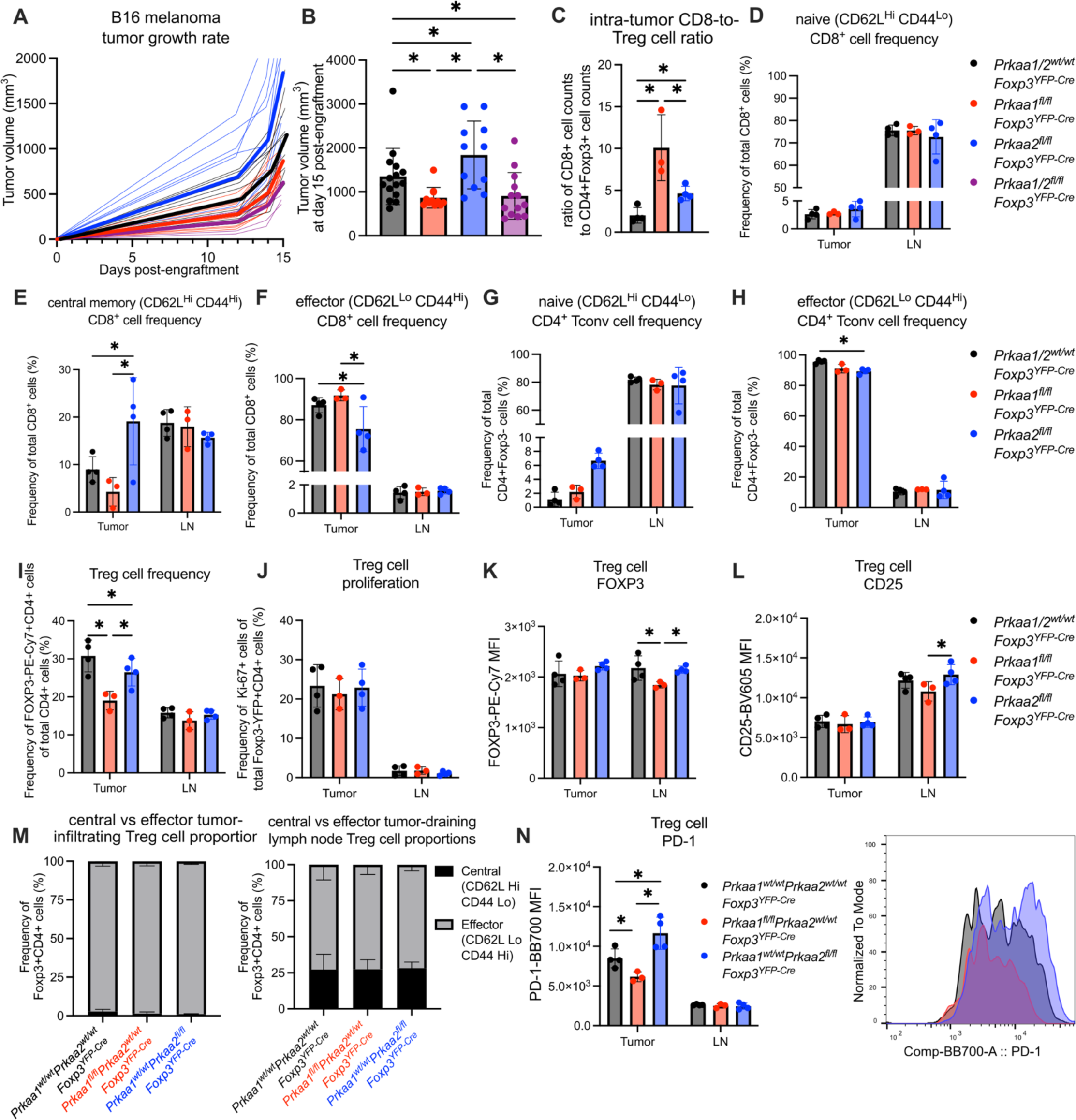
The AMPKα1 and AMPKα2 isoforms have differential contribution to Treg cell suppressive function in the TME. (**A**-**B**) Growth of B16 melanoma tumors (A) and tumor volume at day 15 post-engraftment (B) of *Prkaa1/2^wt/wt^Foxp3^YFP-Cre^* mice (control, *n*=15; includes replicates from Fig 2A-B), *Prkaa1^fl/fl^Foxp3^YFP-Cre^* mice (*n*=11), *Prkaa2^fl/fl^Foxp3^YFP-Cre^*mice (*n*=11), and *Prkaa1/2^fl/fl^Foxp3^YFP-Cre^* mice (*n*=13; includes replicates from Fig 2A-B). (**C**) Ratio of live CD8+ cell counts to live CD4+Foxp3+ cell counts in single cell suspensions of B16 melanoma tumors harvested from the flanks of control (*n*=4), *Prkaa1^fl/fl^Foxp3^YFP-Cre^* mice (*n*=3), and *Prkaa2^fl/fl^Foxp3^YFP-Cre^* mice (*n*=4) at day 15 post-engraftment. (**D**-**F**) of naive (CD62L^Hi^CD44^Lo^; D), central memory (CD62L^Hi^CD44^Hi^; E), and effector (CD62L^Lo^CD44^Hi^; F) CD8 conventional T (Tconv) cells out of total CD8+ cells in tumors and tumor-draining lymph nodes (LN; *n*=4 control, *n*=3 *Prkaa1^fl/fl^Foxp3^YFP-Cre^* mice, *n*=4 *Prkaa2^fl/fl^Foxp3^YFP-Cre^* mice). (**G**-**H**) Frequency of naive (G) and effector (H) CD4+Foxp3-cells out of total CD4+Foxp3-cells in tumors and LN (*n*=4 control, *n*=3 *Prkaa1^fl/fl^Foxp3^YFP-Cre^*mice, *n*=4 *Prkaa2^fl/fl^Foxp3^YFP-Cre^* mice). (**I**) Frequency of CD4+Foxp3+ cells out of total CD4+ cells in tumors and LN (*n*=4 control, *n*=3 *Prkaa1^fl/fl^Foxp3^YFP-Cre^* mice, *n*=4 *Prkaa2^fl/fl^Foxp3^YFP-Cre^* mice). (**J**) Frequency of Ki- 67+CD4+Foxp3+ cells out of total CD4+Foxp3+ cells in tumors and LN (*n*=4 control, *n*=3 *Prkaa1^fl/fl^Foxp3^YFP-Cre^* mice, *n*=4 *Prkaa2^fl/fl^Foxp3^YFP-Cre^* mice). (**K**-**L**) FOXP3-PE-Cy7 (K) and CD25- BV605 (L) mean fluorescence intensity (MFI) of CD4+Foxp3+ cells from tumors and LN (*n*=4 control, *n*=3 *Prkaa1^fl/fl^Foxp3^YFP-Cre^*mice, *n*=4 *Prkaa2^fl/fl^Foxp3^YFP-Cre^* mice). (**M**) Frequency of central (CD62L^Hi^CD44^Lo^) and effector (CD62L^Lo^CD44^Hi^) CD4+Foxp3+ cells out of total CD4+Foxp3+ cells in tumors and LN (*n*=4 control, *n*=3 *Prkaa1^fl/fl^Foxp3^YFP-Cre^* mice, *n*=4 *Prkaa2^fl/fl^Foxp3^YFP-Cre^* mice). (**N**) PD- 1-BB700 MFI of CD4+Foxp3+ cells in tumors and LN (*n*=4 control, *n*=3 *Prkaa1^fl/fl^Foxp3^YFP-Cre^* mice, *n*=4 *Prkaa2^fl/fl^Foxp3^YFP-Cre^* mice). * *q* < 0.05 according to two-way ANOVA of day 15 data from (A) using the two-stage linear step-up procedure of Benjamini, Krieger, and Yekutieli with Q = 5% (**B**). * *q* < 0.05 according to one-way ANOVA using the two-stage linear step-up procedure of Benjamini, Krieger, and Yekutieli with Q = 5% (**C**-**N**). Summary plots show all data points with mean and SD.

**Supplemental Figure 4.**
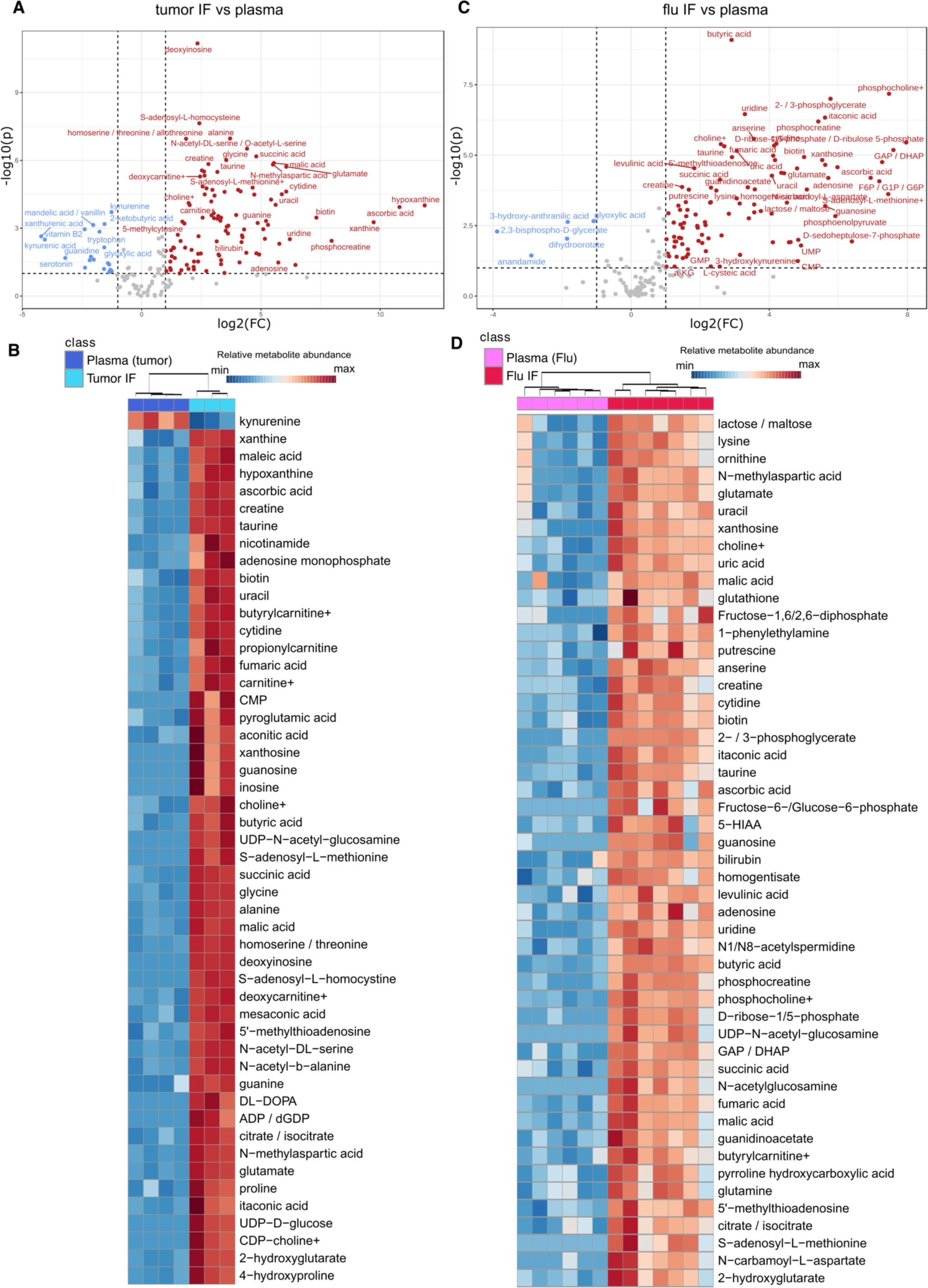
Tumor and influenza virus-infected lung interstitial fluid metabolite abundance compared with plasma. (**A**) Volcano plot of abundance of metabolites detected in tumor (*n*=3) interstitial fluid (IF) and paired plasma (*n*=4). Features with *p* < 0.1 were noted in red if log2(fold-change) ≥ 1.5 or blue if log2(fold-change) ≤ −1.5 when comparing tumor IF versus plasma. (**B**) Heatmap of top 50 differentially represented metabolites between tumor IF versus plasma. (**C**) Volcano plot of abundance of metabolites detected in influenza virus-infected lung (flu, *n*=7) IF and paired plasma (*n*=6). Features with *p* < 0.1 were noted in red if log2(fold-change) ≥ 1.5 or blue if log2(fold-change) ≤ −1.5 when comparing flu IF versus plasma. (**D**) Heatmap of top 50 differentially represented metabolites between flu IF versus plasma.

**Supplemental Figure 5.**
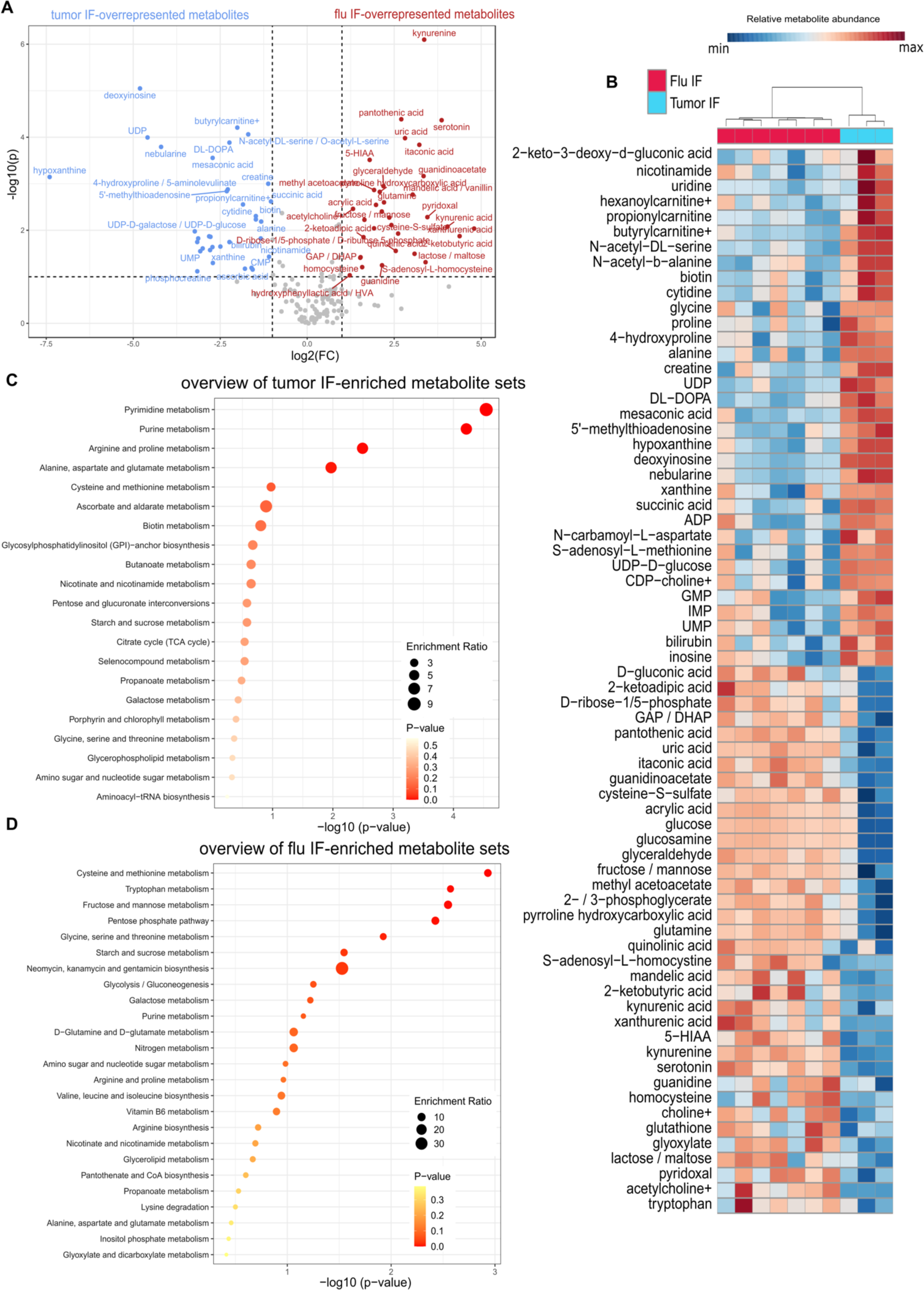
Comparison of tumor interstitial fluid and influenza virus-infected lung interstitial fluid metabolite abundance. (**A**) Volcano plot of abundance of metabolites detected in tumor (*n*=3) interstitial fluid (IF) and influenza virus-infected lung (flu, *n*=7) IF. Features with *p* < 0.1 were noted in red if log2(fold-change) ≥ 1.5 or blue if log2(fold-change) ≤ −1.5 when comparing flu IF versus tumor IF. (**B**) Heatmap of top 50 differentially represented metabolites between flu IF and tumor IF. (**C**) Overrepresentation analysis of significantly (*p* < 0.1) overrepresented metabolites in tumor IF relative to flu IF (**D**) Overrepresentation analysis of significantly (*p* < 0.1) overrepresented metabolites in flu IF relative to tumor IF.

**Supplemental Figure 6.**
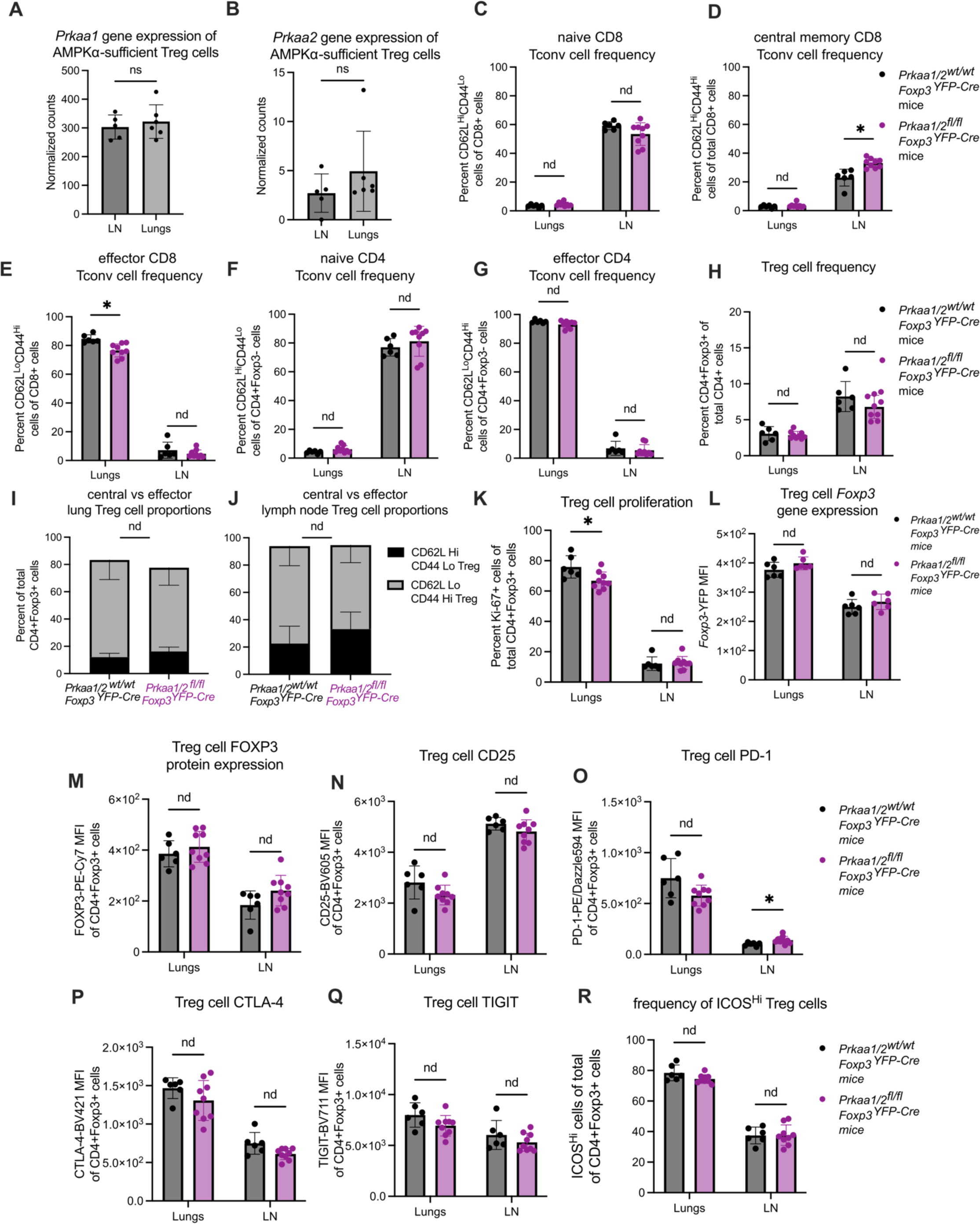
Phenotyping of AMPKα-sufficient and -deficient Treg cells in the lung during viral pneumonia. (**A-B**) Normalized gene expression counts of *Prkaa1* (A) and *Prkaa2* (B) by CD4+Foxp3+ cells sorted from axillary lymph nodes (LN; *n*=5) and lungs (*n*=6) of *Prkaa1/2^wt/wt^Foxp3^YFP-Cre^* (control) mice at day 10 following intra-tracheal inoculation of influenza A/WSN/33 H1N1 (influenza) virus. (**C-E**) Frequency of naive (CD62L^Hi^CD44^Lo^; C), central memory (CD62L^Hi^CD44^Hi^; D), and effector (CD62L^Lo^CD44^Hi^; E) CD8 conventional T (Tconv) cells out of total CD8+ cells in lungs and LN (*n*=6 control, *n*=9 *Prkaa1/2^fl/fl^Foxp3^YFP-Cre^* mice) at day 10 post-influenza virus inoculation. (**F-G**) Frequency of naive (F) and effector (G) CD4 Tconv cells out of total CD4+Foxp3-cells in lungs and LN (*n*=6 control, *n*=9 *Prkaa1/2^fl/fl^Foxp3^YFP-Cre^*mice) at day 10 post-influenza virus inoculation. (**H**) Frequency of CD4+Foxp3+ cells out of total CD4+ cells in lungs and LN (*n*=6 control, *n*=9 *Prkaa1/2^fl/fl^Foxp3^YFP-Cre^* mice) at day 10 post-influenza virus inoculation. (**I**-**J**) Frequency of central (CD62L^Hi^CD44^Lo^) and effector (CD62L^Lo^CD44^Hi^) CD4+Foxp3+ cells out of total CD4+Foxp3+ cells in lungs (I) and LN (J; *n*=6 control, *n*=9 *Prkaa1/2^fl/fl^Foxp3^YFP-Cre^*mice) at day 10 post-influenza virus inoculation. (**K**) Frequency of Ki-67+CD4+Foxp3+ cells out of total CD4+Foxp3+ cells in lungs and LN (*n*=6 control, *n*=9 *Prkaa1/2^fl/fl^Foxp3^YFP-Cre^* mice) at day 10 post-influenza virus inoculation. (**L**) *Foxp3*-YFP mean fluorescence intensity (MFI) of CD4+Foxp3+ cells in lungs and LN (*n*=6 control, *n*=6 *Prkaa1/2^fl/fl^Foxp3^YFP-Cre^* mice) at day 10 post-influenza virus inoculation. (**M**-**Q**) FOXP3-PE-Cy7 (M), CD25-BV605 (N), PD-1-PE/Dazzle594 (O), CTLA-4-BV421 (P) and TIGIT-BV711 (Q) MFI of CD4+Foxp3+ cells from lungs and LN (*n*=6 control, *n*=9 *Prkaa1/2^fl/fl^Foxp3^YFP-Cre^*) at day 10 post-influenza virus inoculation. (**R**) Frequency of ICOS^Hi^ CD4+Foxp3+ cells of total CD4+Foxp3+ cells in lungs and LN (*n*=6 control, *n*=9 *Prkaa1/2^fl/fl^Foxp3^YFP-Cre^*) at day 10 post-influenza virus inoculation. * *q* < 0.05, ns not significant, nd no discovery according to Mann Whitney U test (**A**-**B**) with two-stage linear step-up procedure of Benjamini, Krieger, and Yekutieli with Q = 5% (**C**-**R**). Summary plots show all data points with mean and SD.

**Supplemental Figure 7.**
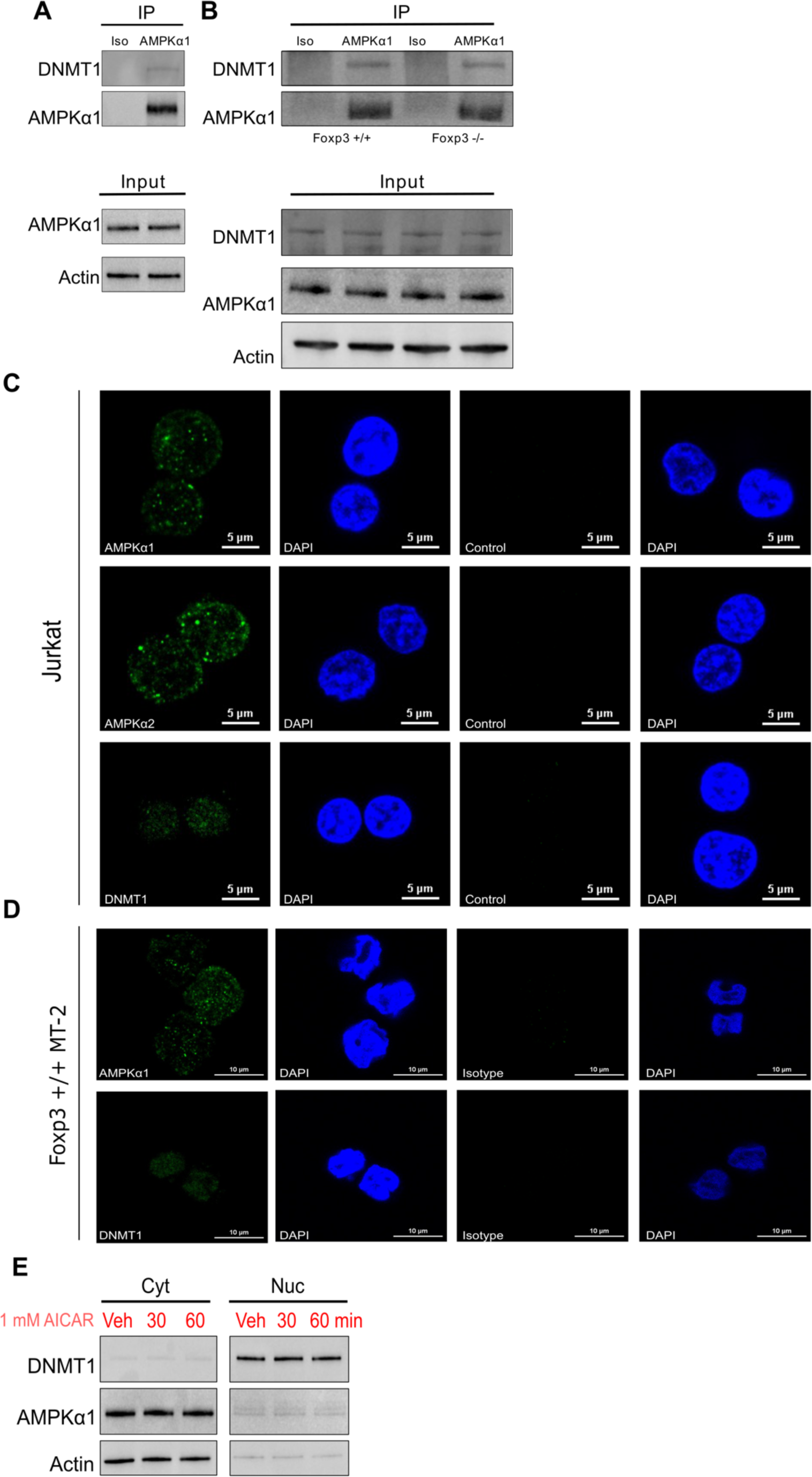
AMPKα1 interacts with DNMT1 in Jurkat and MT-2 cells. (**A**-**B**) anti-AMPKα1 and isotype control immunoprecipitates from Jurkat cell (A) and MT-2 cell (B) lysates blotted for DNMT1 protein. (**C**-**D**) Representative microscopy images of Jurkat cells (C) and MT-2 cells (D) showing AMPKα1 and DNMT1 subcellular localization. (**E**) Immunoblots for DNMT1, AMPKα1, and β- actin on nuclear and cytoplasmic fractions of cell lysates obtained from AMPKα1/α2-sufficient *ex vivo* induced (i)Treg cells treated with either vehicle, 30min AICAR 1mM, or 60min AICAR 1mM.

**Supplemental Table 1.**
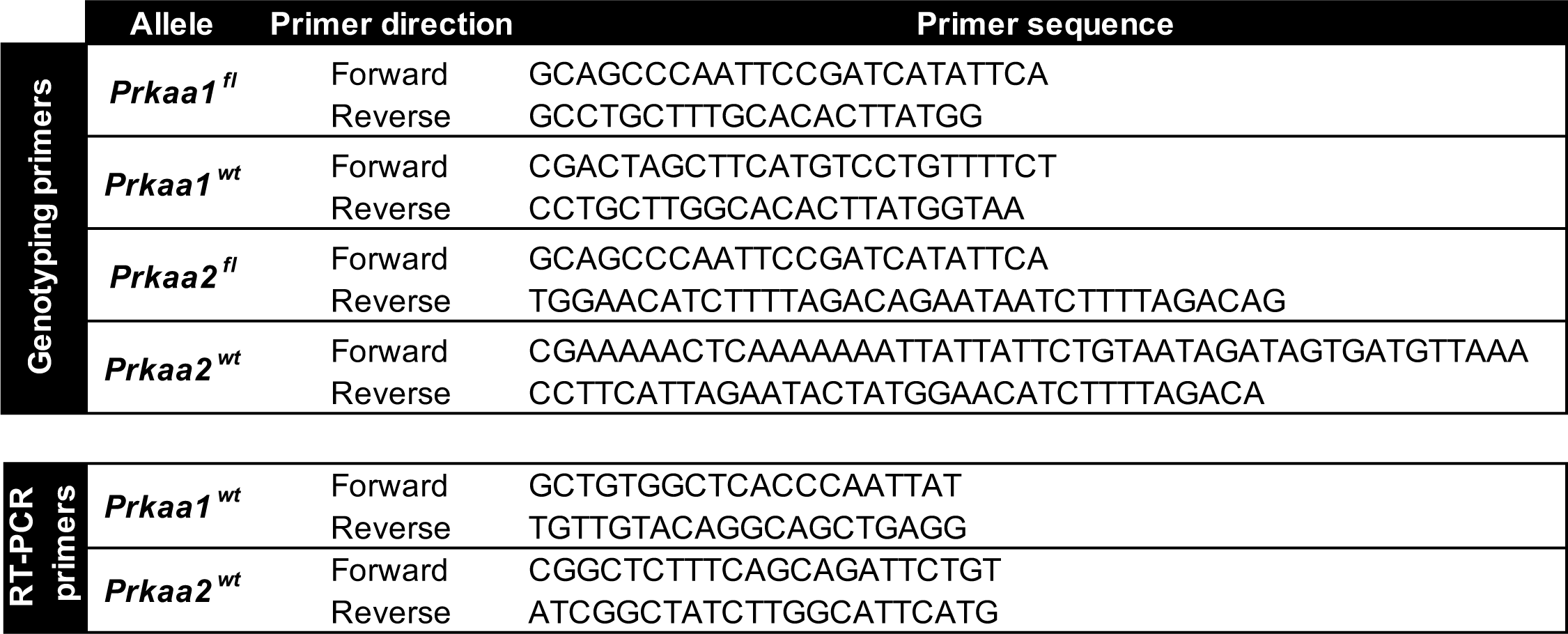
Primer sequences used to genotype the *Prkaa1* and *Prkaa2* loci.

**Supplemental Table 2.** Flow cytometry fluorochromes.

| Antigen/Reagent | Conjugate | Clone | Company | Catalog no. | Notes |
| --- | --- | --- | --- | --- | --- |
| CD45 | BUV563 | 30-F11 | BD Horizon | 612924 |  |
| CD4 | BUV395 | GK1.5 | BD Horizon | 563790 |  |
| CD8 | BUV805 | 53-6.7 | BD Horizon | 612898 |  |
| CD8 | BV711 | 53-6.7 | Biolegend | 100747 |  |
| CD25 | BV605 | PC61 | Biolegend | 102036 |  |
| CD25 | APC | PC61.5 | Invitrogen | 2154040 |  |
| CD62L | APC-e780 | MEL-14 | Invitrogen | 2011195 |  |
| CD44 | BUV737 | IM7 | BD Horizon | 564392 |  |
| PD-1 | PE/Dazzle594 | RMP1-30 | Biolegend | 109116 |  |
| PD-1 | PerCP-eFluor710 | J43 | Invitrogen | 46-9985-82 |  |
| CTLA-4 | BV421 | UC10-4B9 | Biolegend | 106312 |  |
| TIGIT | BV711 | 1G9 | BD Biosciences | 744214 |  |
| ICOS | BV786 | C398.4A | BD Biosciences | 567922 |  |
| Ki-67 | Alexa Fluor 488 | 11F6 | Biolegend | 151204 |  |
| FOXP3 | PE-Cy7 | FJK-16s | Invitrogen | 25-5773-82 |  |
| Fixable Viability Dye eFluor 506 | N/A | N/A | eBioscience | 65-0866-14 |  |
| Absolute counting beads | N/A | N/A | Invitrogen | C36950 | Used according to manufacturer instructions |
| MitoTracker™ Deep Red FM | N/A | N/A | Invitrogen | M22426 | Following surface marker fluorochrome stain, single cell suspensions from harvested tissues were stained with 50 nM Mitotracker Deep Red in complete RPMI for 30 minutes at 37°C. |

