## Supplemental File Legend for "AMP-activated protein kinase is necessary for Treg cell functional adaptation to microenvironmental stress"

**Supplemental File 1.** Peak intensity data of annotated metabolites detected in AMPK-deficient and -sufficient splenic Treg cells of 12–15-week-old mice at homeostasis.

**Supplemental File 2.** Differentially expressed genes detected when comparing AMPK-deficient versus -sufficient splenic Treg cells of 12–15-week-old mice at homeostasis and their corresponding k-means cluster.

**Supplemental File 3.** Differentially expressed genes detected when comparing AMPK-deficient versus -sufficient B16 melanoma tumor-infiltrating Treg cells of 12–15-week-old mice and their corresponding k-means cluster.

**Supplemental File 4.** Peak intensity data of annotated metabolites detected in the interstitial fluid of lungs from influenza virus-infected mice (10 days post-inoculation), the interstitial fluid of B16 melanoma tumors (15 days after subcutaneous engraftment), and paired plasma samples.

**Supplemental File 5.** Peak intensity data of annotated metabolites detected in AMPK-deficient and -sufficient lung Treg cells from 12–15-week-old influenza virus-infected mice (10 days post-inoculation).
